# Fitness and community feedbacks: the two axes that drive long-term invasion impacts

**DOI:** 10.1101/705756

**Authors:** Jean-François Arnoldi, Matthieu Barbier, Ruth Kelly, György Barabás, Andrew L. Jackson

## Abstract

Many facets of ecological theory rely on the analysis of invasion processes, and general approaches exist to understand the early stages of an invasion. However, predicting the long-term transformations of communities following an invasion remains a challenging endeavour. We propose an analytical method that uses community structure and invader dynamical features to predict when these impacts can be large, and show it to be applicable across a wide class of dynamical models. Our approach reveals that short-term invasion success and long-term consequences are two distinct axes of variation controlled by different properties of both invader and resident community. Whether a species can invade is controlled by its invasion fitness, which depends on environmental conditions and direct interactions with resident species. But whether this invasion will cause significant transformations, such as extinctions or a regime shift, depends on a specific measure of indirect feedbacks that may involve the entire resident community. Our approach applies to arbitrarily complex communities, from few competing phenotypes in adaptive dynamics to large nonlinear food webs. It hints at new questions to ask as part of any invasion analysis, and suggests that long-term indirect interactions are key determinants of invasion outcomes.

## Introduction

Predicting the outcome of introducing a new species or phenotype in a resident community is key to answering fundamental ecological and evolutionary questions (Elton 1958). Invasion analysis is invoked to understand adaptation, species coexistence and ecosystem assembly (Law & Morton 1996, Chesson 2000, O’Sullivan *et al.* 2018). It is instrumental in guiding management and conservation efforts in relation to invasive species (Pimm 1991, Williamson 1999, Galiana *et al.* 2014). To analyse the initial stages of an invasion, powerful theoretical approaches exist, built on the notion of invasion fitness or invasibility (Turelli 1978, Metz *et al.* 1995, Geritz *et al.* 1998a, Guo *et al.* 2015, Grainger *et al.* 2019). In essence, these approaches ask under which conditions the invader can grow and spread from a small initial population. The invasibility of a community involves both environmental conditions and direct interactions between the invader and resident species. These properties are the most studied aspects of the invasion process (Jiménez-Valverde *et al.* 2011, Blackburn *et al.* 2011), and determine whether the biotic and abiotic environment is favourable to the invader (Guo *et al.* 2015), and how this can be predicted in terms of the functional traits of the organisms (Eisenhauer *et al.* 2013, MacDougall *et al.* 2009).

But knowing that an invasion can occur in an instantaneous sense, can we predict its long-term consequences (Levine *et al.* 2003), such as whether it drives other species to extinction, or even causes an ecosystem regime shift (Scheffer *et al.* 2001, Gaertner *et al.* 2014, Kotta *et al.* 2018)? These long-term consequences are not only tied to characteristics of the invader and its immediate interaction partners (predators, prey, competitors…), but they can involve the whole web of interactions in the resident community (White *et al.* 2006, Hui & Richardson 2018, Rossberg & Barabás 2019). They may thus be highly unpredictable (Catford *et al.* 2019), both because of their complexity and because few, if any, interaction networks are exhaustively known and accurately quantified (Frost *et al.* 2019). The question is, therefore, whether we can understand the essential features of invader and resident communities that control the qualitative nature of long-term impacts (e.g. benign effect or extinctions) and their order of magnitude.

We propose that classical invasion fitness (Metz *et al.* 1992, Schreiber 2000) plus a novel measure of long-term feedback on the invading population are two complementary dimensions that can be used to characterize the long-term outcomes of invasions. In contrast with invasion fitness, we show that the feedbacks most relevant to longer-term invasion dynamics involve indirect interaction pathways that may run through the entire resident community (White *et al.* 2006). Using invasion fitness and these feedbacks, we then propose a predictor of long-term impacts on the resident community. It is applicable to an arbitrary number of resident species, and to all interaction types, network structures, and functional responses. We show that the invader’s impact can be understood as a change of the community’s biotic environment, and thus, our theory reveals a connection between a community’s robustness against environmental perturbations (Meszéna *et al.* 2006, Ives & Carpenter 2007, Barabás *et al.* 2014) and its vulnerability to invasions.

From our theory it follows that the positioning of invader-community pairs along the two axes of short-term invasion fitness and long-term feedbacks is enough to predict essential features of invasion impacts. We first apply this idea to the simple case of two competitive species, where it naturally connects with known results of coexistence theory and adaptive dynamics (Roughgarden 1983, Metz *et al.* 1996, Dieckmann & Law 1996, Geritz *et al.* 1998b, Champagnat *et al.* 2002, Meszéna 2005, Meszéna *et al.* 2005, Brännström *et al.* 2013). We show, in particular, that when the feedback from the community on the invader is positive, the invasion causes an irreversible shift in community state, signalling the existence of alternative stable states. We then consider linear and nonlinear species-rich model communities, thus illustrating the general predictive power of our theory.

Overall, our work showcases the important information on biotic impacts that can be gleaned from analysing the invader’s dynamics, even without a detailed knowledge of the invaded community. Our approach suggests novel empirical intuitions, revealing indirect interactions and community stability as understudied yet central drivers of invasion dynamics.

## An extension of classic invasion analysis

Invasion analysis (Turelli 1978, Geritz *et al.* 1998a, Williamson 1999, Lewis *et al.* 2016, Barabás *et al.* 2018) typically starts from a stable resident community (comprised of *S* resident species), to which we add a small invading population (species 0). In the following we denote by *N*_0_ the invader abundance and write the resident abundance distribution as a vector **N** = (*N*_*i*_) *i* = 1, …, *S*. We consider general invader-community dynamics of the form

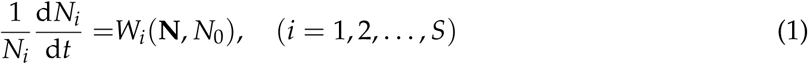

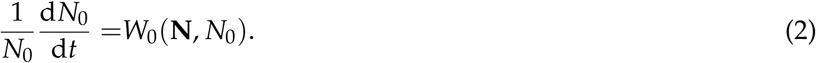

where species growth rates *W*_*i*_ (*i* = 0, 1, …, *S*) are differentiable functions of species abundances. The growth rate *W*_0_ of the invading species, in particular, depends on the abiotic environment and the biotic conditions determined by resident abundances **N**, as well as its own abundance *N*_0_. Assuming that resident species initially coexist with stationary abundances 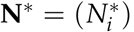, so that *W*_*i*_(**N**^∗^, 0) = 0, the invasion fitness (invader growth rate when rare) is given by

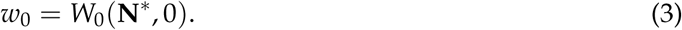

If *w*_0_ < 0, the species cannot grow from rarity. By construction, *w*_0_ describes short-term dynamics immediately after the species’ introduction. It does not account for any feedbacks caused by the invading population itself, since the invader is too rare to have an appreciable influence on the residents. The invasion fitness measures how initially favourable the resident community is to the invader species, both in terms of abiotic and biotic conditions, and common approaches of invasion analysis mainly focus on this quantity (Turelli 1978, Metz *et al.* 1995, Geritz *et al.* 1998b, Williamson 1999, Lewis *et al.* 2016).

Here we propose that long-term effects can be predicted via a simple extension of the invasibility approach. The idea is to compute the feedback experienced by the invading population as it grows and transforms the community. Specifically, we study how the invader’s growth rate *W*_0_ changes as *N*_0_ increases, and modifies the resident species abundances **N**. We claim that by analysing the curve drawn by *W*_0_(**N**, *N*_0_) (thick black in Fig. 2), we can obtain clues on how strongly the invader is transforming the community.

**Figure 1:**
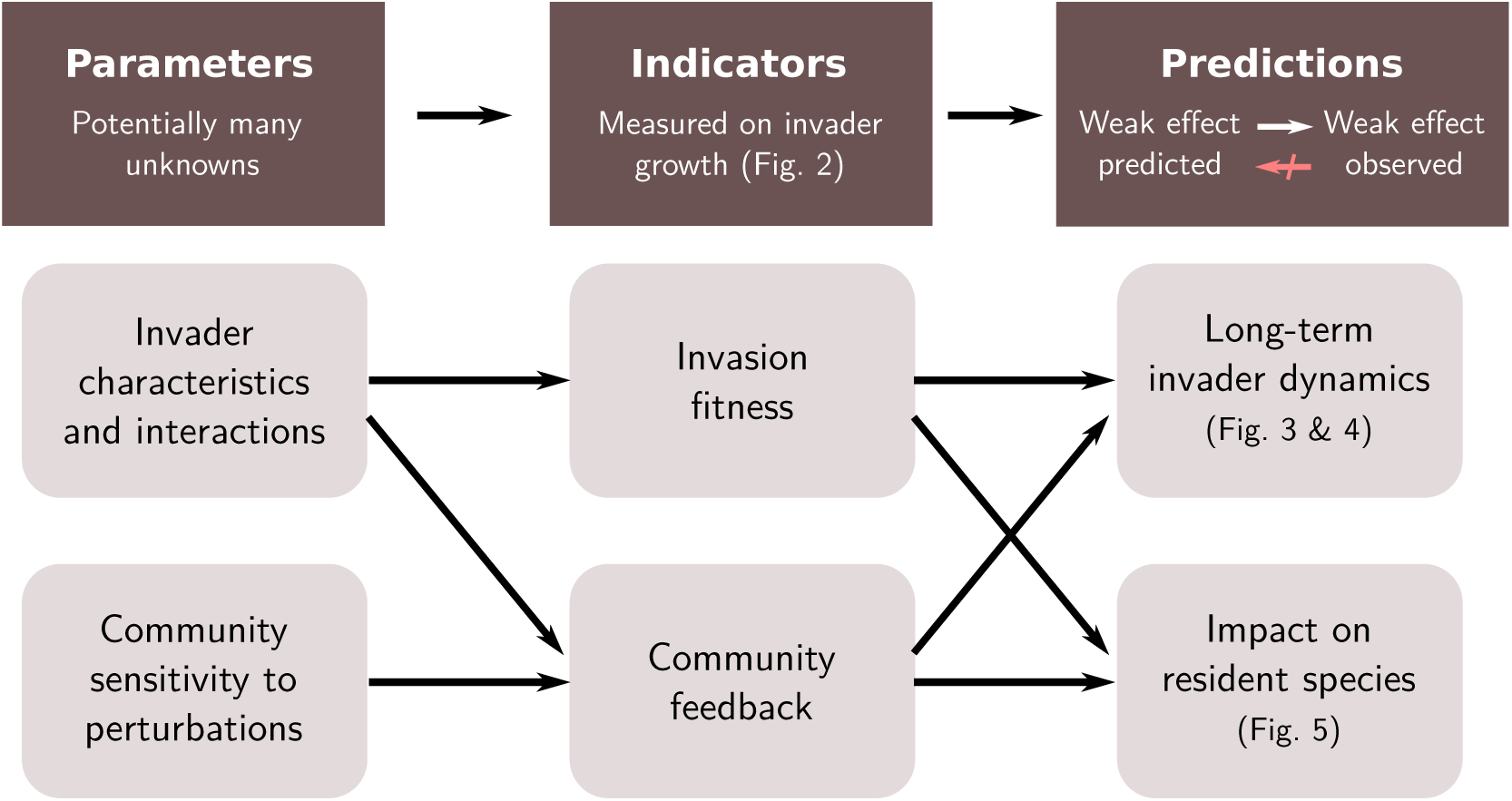
Scheme of our theoretical treatment of long-term invasion dynamics. The invading population’s characteristics and interactions with resident species determines its invasion fitness (growth rate when rare). The invader acts as a perturbation of the resident community. The community’s sensitivity to changes in its biotic and abiotic environment determines its response, and thus the feedback experienced by the invading population as it grows. Invasion fitness and community feedback are features that can be measured from the population growth of the invader, without detailed knowledge of the resident community. Nevertheless, these indicators allow for predictions of long-term outcomes, both for the invader and its impact on the community. While our predictions are extrapolations, they are guaranteed to avoid false negatives: if a weak impact is predicted, a weak impact will be observed.

**Figure 2:**
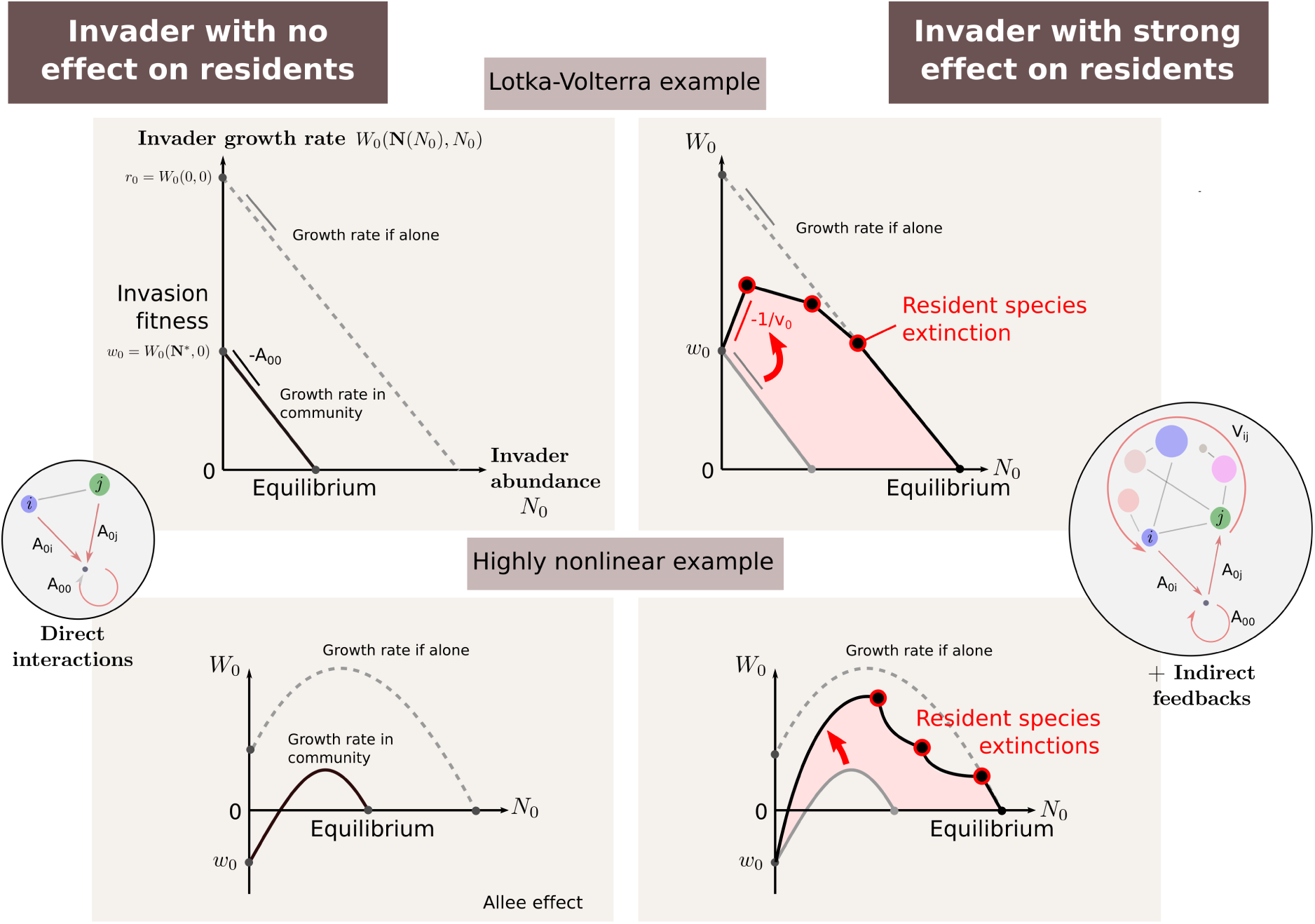
Observing the long-term impacts of an invasion through the invader’s growth curve. In each panel, the invader’s growth rate *W*_0_(**N**, *N*_0_) is plotted against its own abundance *N*_0_. **Left:** Examples where the invader does not influence the resident community. **Right:** Examples where the invader drives resident species extinct. In case all residents disappear, the invader growth curve recovers its shape when alone (dashed lines). In general, we cannot access the whole curve, but only its properties for a small invader population, *N*_0_ ≈ 0. Classic invasion analysis focuses on invasion fitness, *w*_0_ = *W*_0_(**N**∗, 0), the initial growth rate of the invader determined by the resident community state **N**∗. We propose that long-term impacts can be estimated using another local quantity: the initial slope of the invader’s growth rate (see main text). This slope represents how the invader, by transforming its biotic environment, limits itself or accelerates its own growth. We can extrapolate the impact on resident species by comparing this slope to its value −*A*_00_ if residents are not affected (red arrows). **Insets** illustrate that *w*_0_ arises from direct interactions while *v*_0_ encapsulates indirect feedbacks.

Our approach proceeds in two steps, first taking the perspective of the invader, then that of the invaded community. The first step allows us to derive essential characteristics of the invasion dynamics, such as whether the invasion is expected to succeed or fail, and whether it can lead to alternative stable states. In the second step, we explain how, from both the invasion fitness and feedbacks, we can derive a qualitative predictor of long-term impacts on the resident community (e.g., relative change in species abundance, or extinctions). We illustrate the predictive power of our approach on examples that range from a simple linear two-species system to random multispecies Lotka-Volterra communities, and structured non-linear food webs.

### Observing the community through its feedback on the invader

We start from the perspective of the invading population. Our central argument, illustrated in Fig. 2, is that transformations in the resident community are partially reflected in the growth rate of the invader itself. Let us first imagine that we could compute or measure the invader’s density-dependent growth rate in three situations:

a. the true growth rate *W*_0_(**N**(*N*_0_), *N*_0_) as the invader’s abundance *N*_0_ increases, and causes the abundances of resident species to change along a curve **N**(*N*_0_) (black curve in Fig. 2, right panels),
b. the growth rate *W*_0_(**N**^∗^, *N*_0_) observed if the invader had no impact on the community, or the latter was artificially maintained in its original state **N**^∗^ (solid lines, Fig. 2 left panels, thick grey line in right panels),
c. the growth rate *W*_0_(0, *N*_0_) observed if there was no resident community (dashed lines, Fig. 2).

If, for instance, the invader drives all resident species extinct one by one as *N*_0_ increases, then the true growth curve (a) will depart from the impact-less curve (b), and will eventually converge to the curve (c) of the invader alone (Fig. 2 right). More generally, the invader need not cause extinctions, and resident populations could even increase through predation or facilitation. But the fact that curve (a) departs from (b) indicates that the invader is modifying the community. Comparing this deviation to (c) gives a relative measure of the magnitude of this modification.

This simple argument has two caveats. First, we can only measure those changes in the community that feed back on the invader, so we cannot see impacts on resident species that have no direct or indirect effect on *W*_0_. Second, we generally do not expect to be able to fully access these three density-dependent growth curves, in either theory or experiment. This motivates our next step, finding an approximate but readily accessible indicator.

### A local indicator of community feedbacks

We introduce a simple linear indicator of community feedbacks using the invader’s density-dependent growth rate *W*_0_(**N**, *N*_0_), evaluated for *N*_0_ ≈ 0. Despite being defined when the invader is rare, we will show that it allows robust predictions of long-term outcomes, valid beyond the assumption of rarity.

The rarity assumption *N*_0_ ≈ 0 means that the invader acts as a weak additional source of growth or mortality for other species. We define the local effect of species *n* on *m* as *A*_*mn*_ = −∂*W*_*m*_/∂*N*_*n*_, evaluated with the invader at zero abundance and the other species at their resident equilibria. With this notation, the invader induces a perturbation *δW*_*j*_ = −*A*_*j*0_ *N*_0_ on the resident species’ growth rates. In a linear approximation, its effect *δN*_*i*_ on resident abundances is

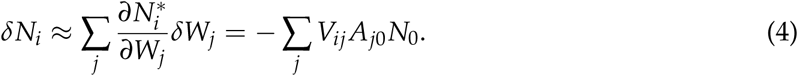

Here 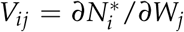 is the long-term sensitivity of species *i* to a change in the growth rate of species *j*. It is computed from the resident interaction matrix *A*_∗_ = (*A*_*ij*_) as 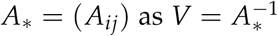 (Levins 1974, Yodzis 1988, Meszéna *et al.* 2006, Aufderheide *et al.* 2013, Barabás *et al.* 2014). The matrix *V*, which we call the *environmental sensitivity matrix*, integrates direct and indirect interaction pathways to determine the community’s sensitivity to changes in species growth rates (Appendix S1). As the invading population affects the abundance of other species, this feeds back on its own growth rate *W*_0_. Under the assumption of weak effects, this represents an incremental change. Using Eq. (4), this change takes the explicit form

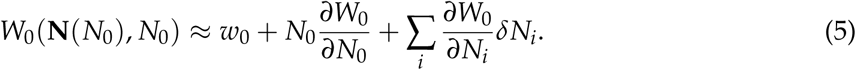

By defining

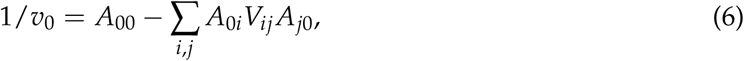

this can be written as

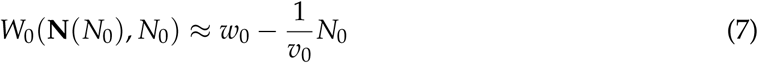

Thus 1/*v*_0_ is the initial slope of the growth rate of the invader. In its explicit expression (6), *A*_00_ represents direct self-regulation of the invader, which can occur even if the community is unaffected by the invasion. The subsequent term encodes indirect effects through changes of the community caused by the invasion.

The difference between 1/*v*_0_ and *A*_00_ is thus a local indicator of community feedback on the invader’s growth rate. By mapping out the relative magnitudes of *w*_0_ and 1/*v*_0_, we identify 5 different possible invasion scenarios. These are described and summarized from the perspective of the invader in Box 1, and justified more formally in Appendix S3. The five scenarios are are: No Invasion (NI), Enhanced Regulatory feedbacks (ER), Reduced Regulatory feedbacks (RR), Positive Feedbacks (PF), and Allee Effect (AE).

#### Box 1

**Predicted scenarios of invasion dynamics (invader perspective)**

Properties of the invader and the resident community determine 5 invasion outcomes that we can map out in a fitness-feedback plane (1/*v*_0_, *w*_0_). We indicate how these scenarios map to regions in that plane, shown by colors in Fig. 3 and 4 (see Appendix S3 for a formal derivation).

**Figure 3:**
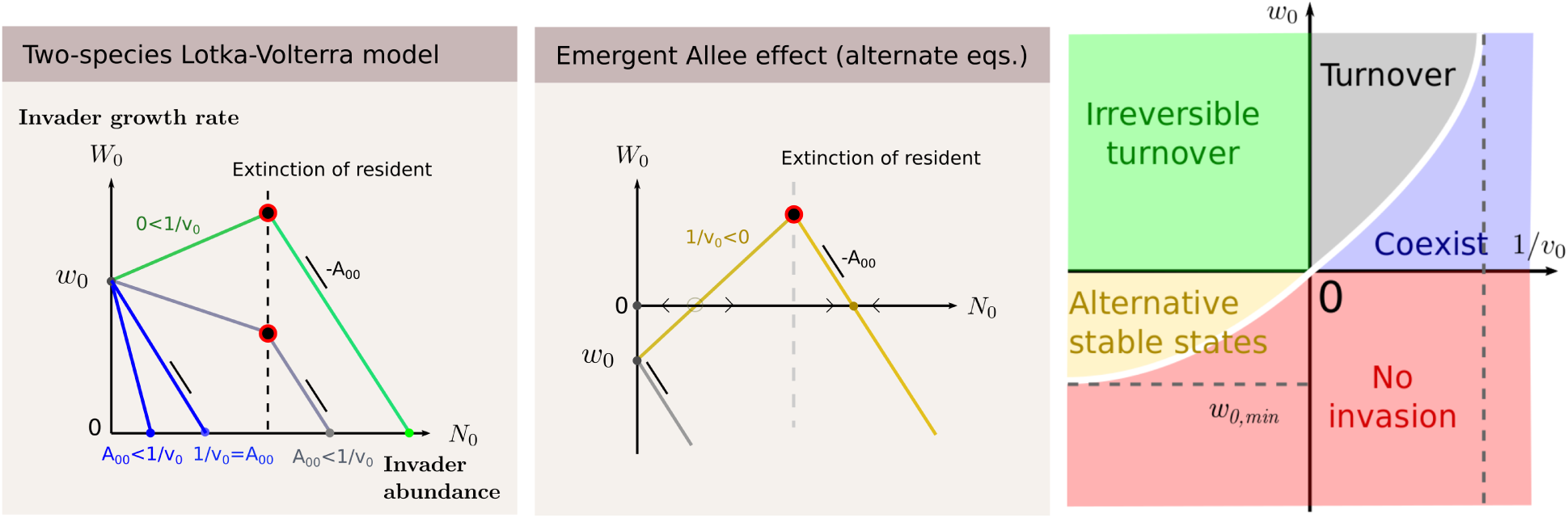
Illustration of the method on a linear two-species competitive system. Left panel: For positive invasion fitness *w*_0_ there are 4 scenarios to consider, determined by the feedback *v*_0_. If 1/*v*_0_ is larger than self-regulation *A*_00_, the invader’s growth is repressed by the resident and cannot displace it (coexistence, in blue also in rightmost panel). If 1/*v*_0_ is positive yet smaller than *A*_00_, the resident species is favouring the growth of the invader, at its expense. Coexistence is still possible but require invasion fitness to be small enough. Otherwise the resident is replaced (reversible turnover, in grey also in rightmost panel). If 1/*v*_0_ is negative, this means that the invader accelerates its own growth, due to the presence of the resident. This diverging feedback loop ends with the exclusion of the resident (irreversible turnover, in green also in rightmost panel). The turnover is irreversible in the sense that there is presence of hysteresis (center panel). Indeed, negative *v*_0_ allows for alternative stable states. In this case, for *w*_0_ ∈ [*w*_0,min_, 0] species 0 will be able to invade only if its initial abundance is high enough (e.g., high propagule pressure is needed for invasion to take place). Furthermore, after a successful invasion reversing abiotic conditions to those pre-invasion will not necessarily allow the resident to re-invade from rarity. The singular case 1/*v*_0_ = 0 (vertical axis in the rightmost panel) is the classical setting of adaptive dynamics, where a mutant phenotype fixes in the population as soon as its fitness is positive.

**Figure 4:**
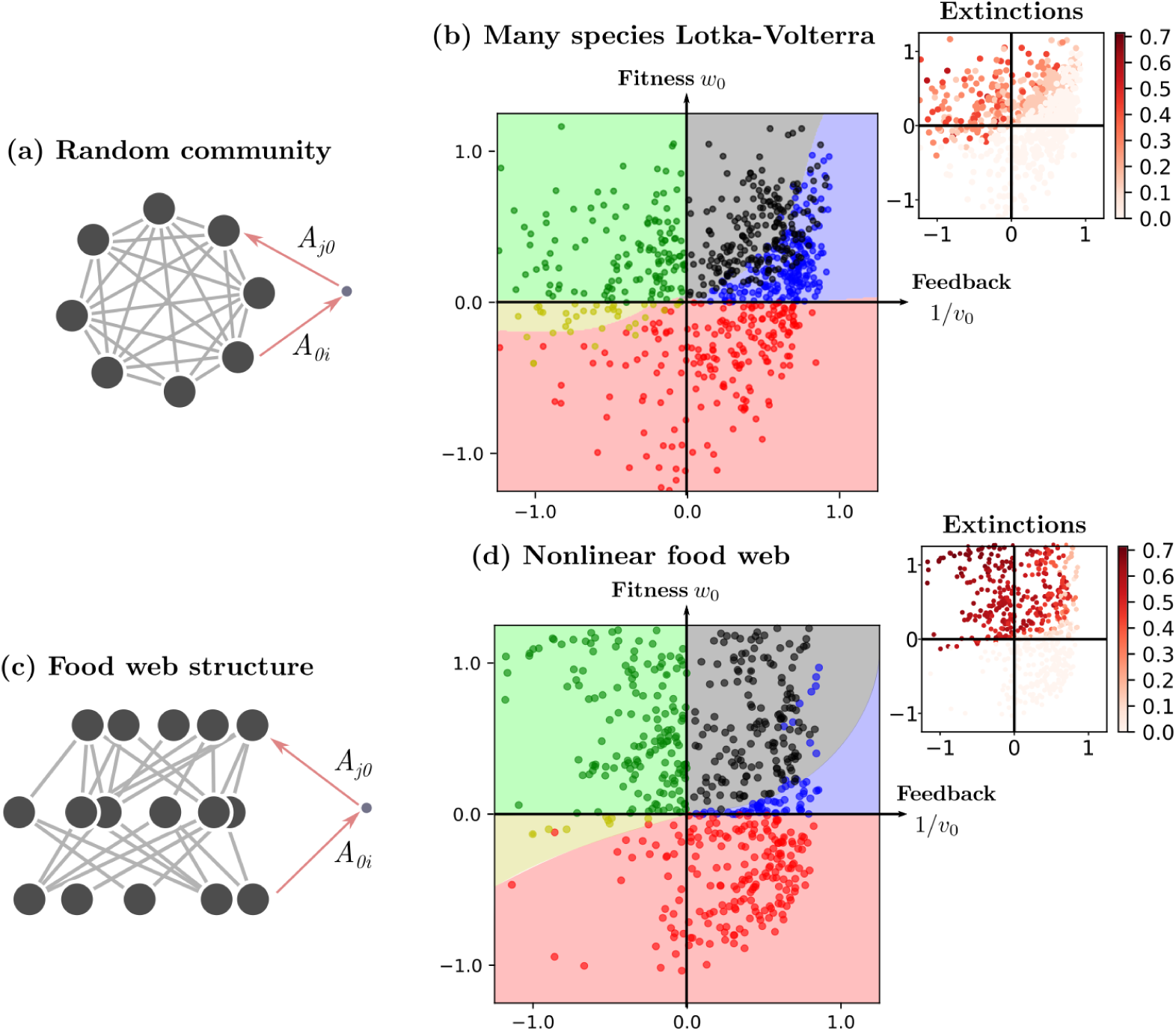
Fitness-feedback maps for **(a-b)** species 0 in a large random Lotka–Volterra community, and **(c-d)** in a two-level food web with nonlinear functional responses (Holling Type 2). The y-axis in maps **(b-d)** is the invasion fitness *w*_0_. The x-axis represents indirect self regulation of the invading species 1/*v*_0_ (Fig. 2). In **(b-d)**, symbols are randomly-drawn invader-community pairs, and we differentiate five outcomes: no invasion (red dots), coexistence (blue dots), turnover (black dots), irreversible turnover (green dots), and alternative stable states where an invader can establish (and cause extinctions) only if its initial population is high enough (high propagule pressure, gold dots). Background colors represent the most frequent outcome at each point of the map, as predicted by a support vector classifier (see Methods). **Insets:** Fraction of resident species going extinct due to the invasion (see Appendix S7 of the Supporting Information for details of the simulation procedure).

NI **No Invasion** (red region). If *w*_0_ < 0, the invasion fails from rarity (but see AE below).

ER **Enhanced Regulation.** If 1/*v*_0_ ≥ *A*_00_ the community only enhances (or does not change) invader self-regulation. Unless *w*_0_ is large, the invasion likely ends with the establishment of a small invader population.

RR **Reduced Regulation.** If 0 < 1/*v*_0_ < *A*_00_ community feedbacks reduce self-regulation. The invader population is modifying its biotic environment to its advantage. The larger *w*_0_ is, the larger this modification.

The two scenarios ER and RR can lead to either coexistence for low invasion fitness *w*_0_ (blue region^*a*^) or replacement of some resident species for large *w*_0_ (black region). In the latter case, if abiotic conditions were modified to lower *w*_0_, the resident species would re-establish, and therefore we call this the *reversible* turnover domain.

PF **Positive Feedbacks** (green region). If 1/*v*_0_ < 0 the response of the community feeds back positively on the invading population. The density-dependent growth rate *W*_0_ first increases with *N*_0_, instead of decreasing towards an equilibrium. This regime displays a tipping point at *w*_0_ = 0 and hysteresis: modifying environmental conditions to increase invasion fitness *w*_0_ above 0 will suddenly allow the invader to establish with finite population, while reducing *w*_0_ back below 0 afterwards will not be enough to eradicate it, as it enters into the AE scenario.

AE **Allee Effect** (yellow region). If 1/*v*_0_ < 0 but *w*_0_ < 0, the invading species cannot grow from rarity, but it may be able to grow from a larger population (Taylor & Hastings 2005).

We finish our local analysis by taking Eq. (7) a step further and noting that invasion ends when 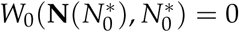. If the invader’s effects are weak enough for the linear approximation Eq. (7) to hold, we get

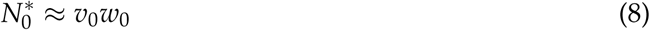

(Appendix S2). As shown below and in Fig. 2 top, this extrapolation of the invader’s equilibrium abundance is exact in any Lotka–Volterra model provided that no resident goes extinct. More generally, it holds in a neighbourhood of *w*_0_ = 0,^1^ as long as *v*_0_ > 0.^2^

### A local indicator of long-term impacts

What matters to us, however, is not the abundance of the invader but its impact on the resident community. By impact, we mean a measure of change of the invaded community. A simple example would be the average relative change in abundance of resident species, which we can estimate using Eq. (4) above. For any species *i* that interacts with the invader (*A*_0*i*_ ≠ 0) we may write

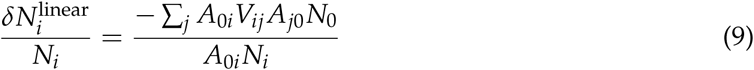

That the numerator bears a close resemblance with the measure community regulatory feedbacks 1/*v*_0_ (Eq. 6) is not a coincidence. In Appendix S1 we derive an expression for 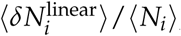, the mean abundance change among species that interact with the invader, divided by the mean pre-invasion abundance of those same species (see Box 2). Remarkably, its expression (Eq. 10 in Box 2) depends solely on local invader characteristics: its relative invasion fitness and regulatory feedback. The impact extrapolation 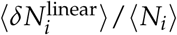 is not intended to be an accurate, quantitative measure of the relative change in resident abundance caused by the invasion. It is a proxy, that can be used to make qualitative predictions of the actual impact (i.e some measure of change in the resident community). In fact, if the proxy is large, it is unlikely to give an accurate prediction of abundance change, because nonlinear effects will likely play out (e.g., extinctions). Although it is possible that we overestimate actual impact (possible false positive), if the proxy is small, it will be accurate (because the dynamics will be linear) and there will thus be a weak impact (no false negatives). Of course, in practice (e.g. in an experimental setting), the uncertainty of such predictions will be inevitably inflated by additional sources of error.

#### Box 2

**Predicted impacts of invasion (community perspective)**

From a local analysis of invasion fitness and feedbacks we deduce a indicator of impacts on the community. We claim that this indicator leads to predictions that do not make false negatives. This indicator reads (see Fig. 5 for an illustration and Appendix S1 for a derivation):

**Figure 5:**
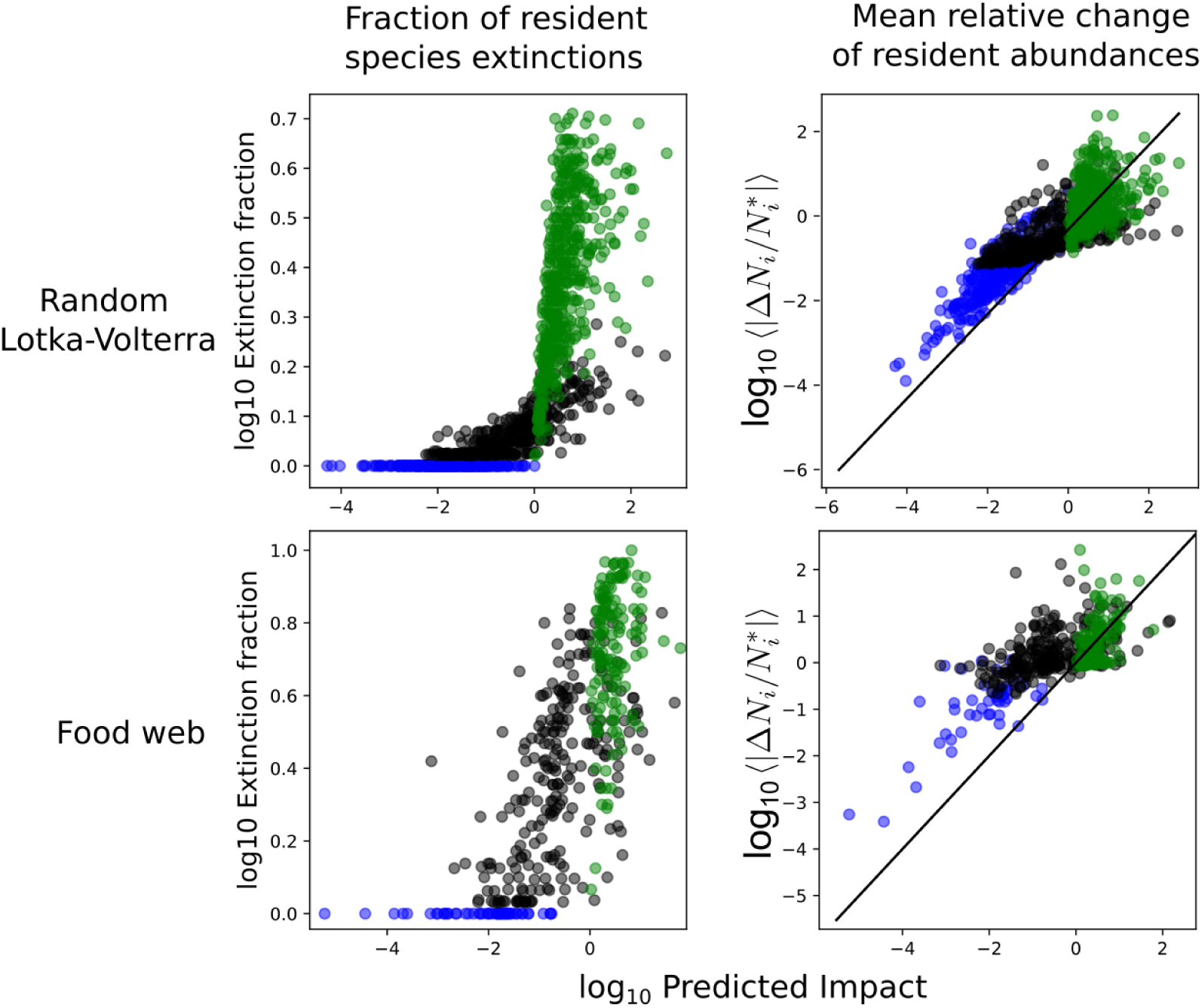
Impact prediction based on the linear extrapolation of impact (10) vs two actual outcomes of simulated invasions: **(left)** fraction of extinctions and **(right)** mean change of relative abundance 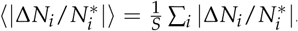. As in Fig. 4, we consider random Lotka–Volterra communities **(top)**, and two-level food-webs with nonlinear functional responses (Holling Type 2, **down**). Dots correspond to randomly-drawn invader-community pairs, and colors are associated with the domain in the fitness-feedback maps of Fig. (4). **Blue**: no extinctions, **black**: reversible turnover, **green**: irreversible turnover. Spearman rank-order correlation coefficient *ρ* between predicted impact and fraction of extinction is *ρ* = 0.75 for Lotka–Volterra communities, and *ρ* = 0.57 for the non-linear food-web ensemble. Correlation coefficient between predicted impact and mean abundance change is *ρ* = 0.60 for the Lotka–Volterra communities, and *ρ* = 0.44 for the non-linear food-web ensemble. See Appendix S7 for details of the simulation procedure.

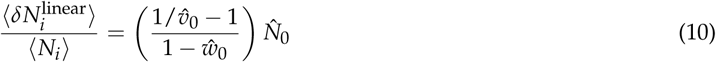

where

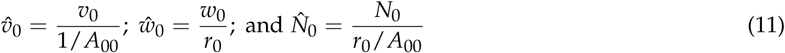

1/*A*_00_ is the feedback that the invader would experience *if the invader had no effect on the community* (slopes on left panel of Fig. 2). This makes 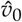 a measure of relative regulatory feedbacks. Similarly, *r*_0_ = *W*_0_(0, 0) is the invasion fitness *if the community had no effect on the invader* (intercept of the dashed lines in Fig. 2). This makes 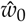 a measure of relative invasion fitness.

The value chosen for 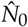 (a generalized notion of relative yield) depends on the scenario of invasion listed in Box 1.

NI: the invasion fails so that 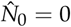

ER & RR: Based on (8), we estimate 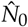 as 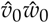.

PF: The invader abundance surges to unknown values. We conservatively set 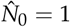.

AE: From rarity, the invasion fails. If the initial abundance is large enough, the invader abundance surges to unknown values. In simulations (Fig. 5) we only consider the former, thus 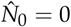.

## Applications

### Linear two-species community

A species invading a resident population of a competing species provides an enlightening illustration of the above reasoning. In this elementary example, our approach naturally connects with known results of competition theory (Roughgarden 1983, Tilman 1982, Meszéna *et al.* 2006) and adaptive dynamics (Metz *et al.* 1992, 1995, Geritz *et al.* 1998b, Meszéna 2005, Meszéna *et al.* 2005, Brännström *et al.* 2013).

Consider a competitive Lotka–Volterra model, so that the growth rate of the invading species takes the linear form

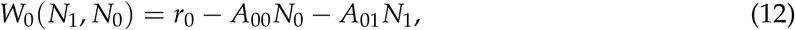

Here *r*_0_ = *W*_0_(0, 0) > 0 and all interaction terms *A*_*ij*_ are positive constants. If the resident population is initially at carrying capacity *K*_1_, the invasion fitness (3) of species 0 is

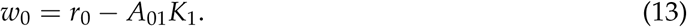

The sensitivity matrix of the resident is a single number 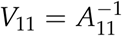 and

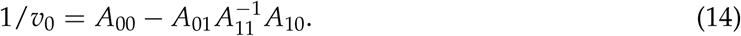

Because growth rates are linear functions of abundance, our expression (7) is exact (this is true for any Lotka–Volterra system with arbitrarily many species):

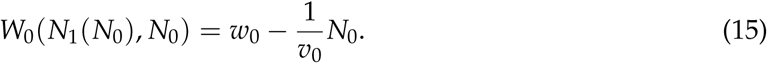

And, therefore, if species coexist, we have that 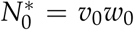. We thus see that *v*_0_ represents the invader’s long-term sensitivity to a change in its abiotic environment (e.g changing *r*_0_) *in the presence of the resident species*.

In this linear example, our proxy of impact, Eq. (9), is exact in the coexistence region (and when the invasion fails) and will underestimate impact if the resident goes to extinction. Let us focus on the invasion scenarios listed in Box 1. We show in Appendix S3 that the x-y axes and one curve, respectively:

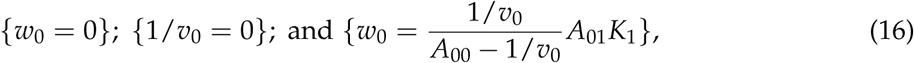

together delineate the 5 regions in the fitness-feedback plane shown in Fig. 3.

The invasion outcomes follow the expectations listed in Box 1, but we can make a few useful remarks. For Lotka–Volterra dynamics, there cannot be coexistence if 1/*v*_0_ < 0: the growth rate *W*_0_ will keep growing with invader abundance *N*_0_ until it causes the extinction of the resident species (Fig. 3 left). If *w*_0_ < 0, the invader cannot grow from rare, but it may be able to grow from finite abundance. Hence, the feedback through the resident species can create an Allee effect on the invader (Fig. 3 center). The state reached (invader or resident) then depends on initial conditions (yellow region in Fig. 3 right).

The singular case 1/*v*_0_ = 0 (y-axis) suggests perfect replacement: an invader with *w*_0_ > 0 will replace a resident. Here a direct connection with adaptive dynamics theory (Geritz *et al.* 1998b, Champagnat *et al.* 2002, Meszéna 2005, Brännström *et al.* 2013) can be made if the invading population has a mutant phenotype, differing only slightly from the resident population. This means that intra- and interspecific interaction strengths are almost exactly equal. By Eq. (14), this implies 1/*v*_0_ ≈ 0. In this case a coexistence state can only exist under special circumstances (a branching point, see Appendix S5). In general, however, as soon as the mutant can invade, it fixes in the community and drives the resident population extinct.

### Multi-species communities

We illustrate in Fig. 4 how the results of the two species case are representative of the behaviour of much more complex models. We generated many-species communities with random Lotka– Volterra interactions and two-level food-webs with nonlinear (Holling Type 2) functional responses. For each communities, we simulated population dynamics until an equilibrium was reached **N**^∗^. We then generated hundreds of invaders with randomly distributed interactions (see Appendix S5), and simulated the outcome of the invasion of each invader separately, monitoring the fraction of species extinctions, their change in abundance, but also checking whether the outcome depended on the initial invader abundance (checking for multi-stability). We see in Fig. 4 that the five possible regions (no invasion, coexistence, turnover, irreversible turnover, and alternative stable states) identified in two-species case are still well ordered in the fitness feedback plane (1/*v*_0_, *w*_0_). In fact, we identify in Appendix S3 the boundaries in the fitness-feedback plane that separate these five outcomes.

In Fig. 5 we compare actual measures of invasion impact – fraction of resident extinctions and mean change of relative abundances– to the linear extrapolation of impact (10) based solely on relative fitness and feedback (Box 2). We see a good correlation of both metrics with our predictor. In the coexistence region (blue dots), resident species do not go extinct, but this does not mean that the invader has no impact – in fact, it may be precisely because of the impact on resident abundances that the invader’s growth is limited (ER regime in Box 1).

Overall, we see that invaders with higher fitness and weaker regulatory feedbacks will have larger impacts: in particular, in the Positive Feedback (PF) regime of Box 1 (green dots), we see that many residents go extinct after invasion, while others might have their abundances increase dramatically^3^. These highly non-linear effects are poorly predicted by our linear extrapolation, yet –as expected– the latter makes no false negatives: we do not observe high impacts when low impacts are predicted.

## Discussion

The invasion dynamics of a species or phenotype can be a highly complex process, inducing significant transformations of the resident community. Classic invasion analysis mainly focuses on the initial stages of invasion, when a species is establishing in a resident community, and either grows and spreads, or goes extinct (Williamson 1999, Blackburn *et al.* 2011, Metz *et al.* 1992). For theory (Lewis *et al.* 2016), this perspective allows a local analysis: one only needs to know the biotic and abiotic environment that the invader perceives at the time of introduction to predict whether the invasion will be successful.

Our contribution is to show that a similar local analysis can shed light on long-term impacts, and assess whether a successful invasion will act only as a slight perturbation of the community, or cause large shifts. While predicting the exact outcome of the invasion is generally out of reach, we can determine whether this outcome is likely to strongly alter the pre-existing community.

We revealed that, together with the classic notion of invasion fitness *w*_0_, the analysis of another quantity, *v*_0_, allows a qualitative prediction of long-term invasion outcomes. *v*_0_ encapsulates the indirect feedbacks experienced by the invading population, as it expands and impacts the rest of the community. Negative feedbacks ensure that the invader will be quickly contained, and will typically coexist at moderate abundance with resident species. Positive feedbacks signal that the invader’s growth will accelerate as the community undergoes significant transformations, such as species extinctions, until a qualitatively different equilibrium is reached. In the latter case, an invader with a weak (or even negative) fitness advantage can achieve disproportionate impacts in the long term on resident species, especially if its initial abundance is high.

A parallel can be drawn with studies of ecosystem engineers –organisms that alter the abiotic features of ecosystems (Wright & Jones 2006). The population growth of an ecosystem engineer is subject to a feedback from the modified abiotic environment (Cuddington *et al.* 2009). Here we have transposed this idea, revealing the critical role in invasion dynamics of feedbacks from the *biotic* environment.

### Alternative stable states in high-diversity communities

Our analysis is a step towards understanding when alternative stable species compositions can emerge from community dynamics (Figs. 3-4). Alternative stable states have mostly been discussed in the context of low-dimensional models of ecosystem functioning and regime shifts (Scheffer *et al.* 2001). But many ecological phenomena hinge on the existence of alternative states in complex, high-dimensional communities (Gilpin & Case 1976, Dakos 2018, Bunin 2018). Priority effects are a common feature of community assembly, and imply that different stable compositions can become established, depending on initial biotic conditions (Law & Morton 1996, Fukami & Nakajima 2011). Sharp spatial boundaries (ecotones) can arise between alternate communities in a homogeneous or smooth environment (Liautaud *et al.* 2019), and a perturbation can push a community from one state to the other.

From our work, it follows that those phenomena ought to be deeply entangled with the communities’ long-term response to invasions (Gaertner *et al.* 2014, Kotta *et al.* 2018). This connection is strongest when there is at most one equilibrium per species composition. Alternative stable states are then necessarily associated with different compositions, and shifts between states can be triggered by invasions. In this case, a compositional shift may or may not imply an ecosystem regime shift (since species can be functionally redundant) but any regime shift must involve species invasions and extinctions.

Conversely, the alternative stable state perspective on invasions allows us to understand two essential phenomena which are not readily captured by classic invasion analysis. When there are positive feedbacks on the invader via the community (1/*v*_0_ < 0) a minute change in environmental conditions can have a disproportionate effect on the community, if it allows the invader to reach a positive invasion fitness. In other words, the invasion threshold *w*_0_ = 0 is, in this case, a *tipping point* (Scheffer *et al.* 2001). Furthermore, positive feedbacks create an emergent Allee effect, allowing a species to persist at a large abundance even if it would not be able to invade from rarity (Taylor & Hastings 2005). We can thus identify when invasion success depends on invader abundance, whether considered as an initial population (priority effect) or an influx from the outside (propagule pressure).

### Invader-as-perturbation and stability

Our analysis shows that the invader can be treated as an environmental press perturbation. This perturbation causes a response from the invaded community, which will feed back on the invader’s growth, thereby changing the strength of the perturbation that the invader represents, and so on.

This perspective on invasion dynamics thus brings together two important facets of ecological stability (Donohue *et al.* 2016): resistance to environmental change and long-term response to invasions. A resident community that is more sensitive to environmental change is also more likely to allow invading species to experience positive indirect feedback. From our formalism we can make this statement more precise, and further clarify how invasion dynamics intertwine the concepts of perturbation, and response to a perturbation. This is particularly clear when considering the *worst-case* bound for regulatory feedbacks 1/*v*_0_:

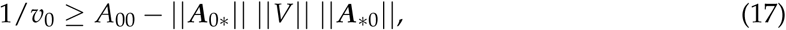

where ‖ · ‖ is the Euclidean norm of interaction vectors ***A***_0∗_ = (*A*_0*j*_), ***A***_∗0_ = (*A*_*i*0_). At first sight, the contribution of the interactions between invader and residents is straightforward: the stronger the interactions, the larger the norms ‖***A***_0∗_‖ and ‖***A***_∗0_‖. If they have the right sign, they can reduce regulatory feedbacks and even allow a phase of runaway growth (if 1/*v*_0_ < 0 cf. Box 1). But the worst-case expression also involves the norm^4^ ‖*V*‖ of the environmental sensitivity matrix. The latter measures the community’s maximal amplification of an environmental press perturbation (Arnoldi *et al.* 2016).

Thus, the largest feedback is generated when invader-community interactions ***A***_0∗_ align with the direction of *the worst-case perturbation* of this community; and when the community-invader interactions ***A***_∗0_ align with the associated direction of *largest response*. This will feedback on the invader’s growth, thereby changing the strength of the perturbation that the invader imposes on the community. In other words, invasions should not only be seen as mere external perturbations, but as perturbations that are shaped by the response that they elicit.

Given a community, the above reasoning provides a method to find its most potent invader^5^ (see Appendix S6). More importantly it showcases a formal link between stability in the face of environmental perturbations (represented by ‖*V*‖), and the long-term impacts of biological invasions (via *v*_0_). This reasoning also clarifies some differences. In particular, if stability relates to our measure of feedback *v*_0_, the latter is only one of the two important drivers of invasion impacts, the other being invasion fitness *w*_0_.

### Empirical connections

Our theoretical analysis yields multiple potential empirical connections, for understanding and predicting outcomes for the invader or the resident community, or using invaders as probes of the community’s stability and response.

We started by a general argument for estimating invasion impacts on a community by looking only at changes in the invader’s density-dependent growth rate, even if we cannot access any information on the community itself. This argument could, in principle, be tested experimentally by reconstructing the three fitness curves of Fig. 2. This would require to observe the population dynamics of the invading species in different contexts (alone or at least in a known biotic context and in the community of interest), thus suggesting a comparative approach to invasion dynamics. An important caveat and potential source of misunderstanding, however, is that these fitness curves should represent long-term effects: an experimental test would involve maintaining a fixed invader abundance *N*_0_, then letting the community settle in a new state **N**(*N*_0_), before measuring the resulting growth rate *W*_0_, and repeating this process for different values *N*_0_. The value of such test would be to reveal whether we can truly get accurate estimates of impact on resident communities through their fingerprint in the dynamics of the invader itself.

We also proposed less systematic but more accessible indicators of community response. As noted above, our analysis relates invasion outcomes to the sensitivity of the resident community to environmental changes. It is however essential to remark that we define community sensitivity as the degree to which species interactions amplify or attenuate the reaction of individual species. It may be that certain ecosystems appear more sensitive because each species, on its own, reacts more strongly to the environmental change; and we do not expect this to relate to invasion impacts. Therefore, to empirically validate our prediction, we will have to carefully tease apart the component of stability that is due to species interactions. This being done, we predict that invasions should have greater impacts in ecosystems that are less stable to environmental changes, and conversely, that response to an invasion could be used a a probe to predict the effect of other perturbations. In any case, our theoretical work demonstrates that long-term feedbacks must, one way or another, be estimated in order to predict invasion outcomes.

The fact that resident species can create density dependence on invaders, with consequences such as Allee effects, tipping points and hysteresis, has been discussed in various theoretical and empirical contexts (Courchamp *et al.* 1999, 2008, Kramer & Drake 2010, Anic *et al.* 2015). In most cases, these discussions have focused on a small number of resident species, in direct interaction with the invader. Our work proposes that these intuitions can be extended to many-species communities. Our theory also suggests that the nature of interactions can play an important role on invasion outcomes: we noted in our two-species analysis that asymmetrical (e.g. predator-prey) interactions will generally lead to negative feedbacks, meaning that they are less likely to give rise to emergent Allee effects; however, positive feedbacks between predator and prey could still arise, e.g. due to indirect effects through competitors McLellan *et al.* (2010), which our many-species analysis would capture.

### Conclusion

Our work underlines the potential for cross-fertilization between the literature on ecological community stability, and the many ecological and evolutionary approaches based on invasion analysis. These approaches, such as adaptive dynamics (Metz *et al.* 1995), have often focused on few-species models and short-term outcomes. Our results suggest that extensions can be made toward many-species networks and long-term feedbacks. Important qualitative properties, such as the possibility of alternate states (McNally & Jackson 2013), or evolutionary branching (Geritz *et al.* 1998a, Doebeli & Dieckmann 2000, Champagnat *et al.* 2002)), could be within reach of a general approach in complex communities.

This general method lends itself to analyzing particular theoretical or empirical interaction networks. Studying diverse systems, from food webs to competitive guilds or complex microbial communities, could help test our predictions about which features contribute more to short-term invasion success or long-term impacts. Our work constitutes a promising avenue to refine predictions on the role of other large-scale properties of an interaction network, such as conectance, directedness, nestedness or modularity (Lurgi *et al.* 2014, Barbier *et al.* 2018), on invasion vulnerability. While we interpreted our results in the context of ecological dynamics, this method is readily extended to other biological dynamics, and could shed light on complex evolutionary dynamics in the presence of phenotypic diversity (Venkateswaran & Gokhale 2019, Kotil & Vetsigian 2018).

## Acknowledgments

We thank Bart Haegeman for helpful comments on a previous version of this manuscript. JFA and ALJ were supported by an Irish Research Council Laureate Award IRCLA/2017/186. MB was supported by the TULIP Laboratory of Excellence (ANR-10-LABX-41) and by the BIOSTASES Advanced Grant, funded by the European Research Council under the European Union’s Horizon 2020 research and innovation programme (666971).

### S1 A proxy to estimate long-term impacts

For any species *i* that interacts with the invader (such that *A*_0*i*_ ≠ 0) we may write

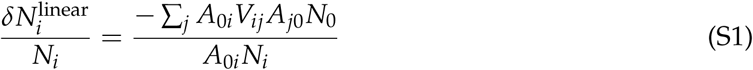

Notice in the numerator the close resemblance with our local expression of community regulatory feedbacks (6) from main text. The sum over species of the numerator is

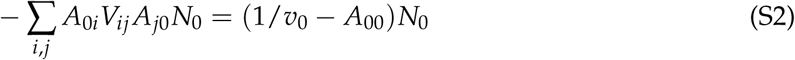

Note that 1/*A*_00_ is the feedback that the invader would experience *if the invader had no effect on the community* (slopes on left panel of Fig. 2 in main text). The sum over species of the denominator in (S1) is

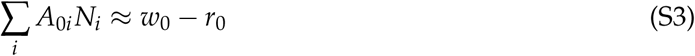

where *r*_0_ = *W*_0_(0, 0). By symmetry with the feedback term, *r*_0_ is the invasion fitness *if the community had no effect on the invader* (intercept of the dashed lines in Fig. 2 in main text). Thus the mean abundance change among species that interact with the invader, divided by the mean pre-invasion abundance of those same species (see Box 2).

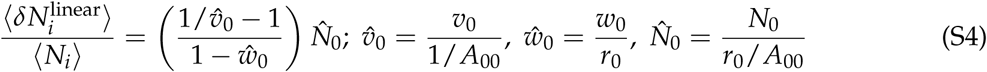

This gives us an approximate, linear extrapolation for average relative change of resident species abundances. We see that it depends two non-dimensional features of the invading population: its relative invasion fitness 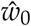, and the relative strength of feedback 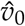. Furthermore, the estimate for the value of 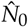 in (S4) will depend on the signs of fitness and feedback. If *w*_0_ < 0 there is no invasion, 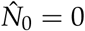 (thus no impact). If both *w*_0_ and *v*_0_ are positive then we can use the coexistence approximation (6), so that 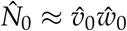. Finally if *v*_0_ < 0, we know that the invader abundance will be large no matter how small its invasion fitness. In this case, a conservative estimate of impact can be deduced by simply setting 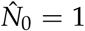. This impact extrapolation (S4) is not intended to be an accurate, quantitative measure of the relative change in resident abundance caused by the invasion. It is a proxy, that can be used to make qualitative predictions of the actual impact (i.e some measure of change in the resident community). In fact, if the proxy is large, it is unlikely to give an accurate prediction of relative abundance change, because non linear effects will likely play out (e.g extinctions). Although it is possible that we overestimate actual impact (possible false positive), if the proxy is small, it will be accurate and there will thus be a weak impact (no false negatives).

### S2 General expression of invader abundance in the coexistence state

Consider a general community model of the form

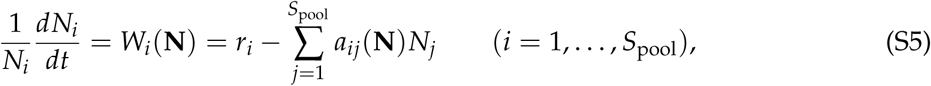

where *r*_*i*_ denotes species’ intrinsic growth (or mortality) rates, which are negative for strict consumers. The terms *a*_*ij*_ could encode any type of inter-specific interaction, and any functional response. The sign convention chosen is that a positive *a*_*ij*_ would be an antagonistic interaction (e.g., competition or predation). The diagonal terms *a*_*ii*_ denote self-interactions (e.g self-regulation when positive). However, we do not assume that there are direct self-interaction terms for all species. In food web models, for instance, one rarely assumes intra-specific interactions for predators (e.g., cannibalism). Before performing the invasion analysis, we do assume that the resident community has reached an equilibrium 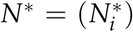 comprised of *S* ≤ *S*_pool_ persisting species. For conciseness we will write 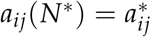. If a new species is introduced with a small invasion fitness

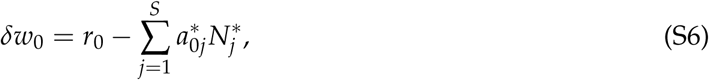

it will grow form rarity. We assume that it eventually reaches a low abundance *δN*_0_. Under this assumption, resident species abundances must have consequently changed by a small amount *δN*_*i*_ to satisfy

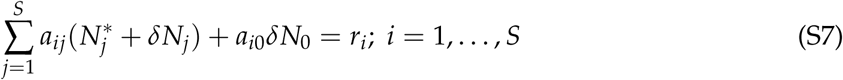

to first order in *δN*_0_ and *δN*_*i*_. This expression implies that

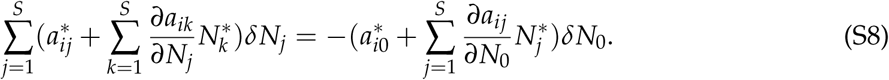

We now define the effective resident interaction matrix *A* = (*A*_*ij*_) as

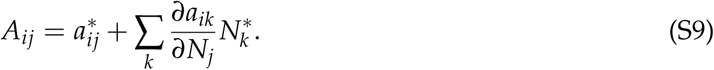

*A*_*ij*_ encodes both direct interactions between *j* and *i* as well as the effect that species *j* can have on other interactions affecting species *i* (e.g., higher order interactions). Also, this term encodes effective self-interactions, that can be non-zero even if *a*_*ii*_ = 0 (e.g., predator interference). Furthermore, we can directly generalize the above expression of *A*_*ij*_ to include effective interactions *A*_*i*0_ between invader and resident species. With these notations we get

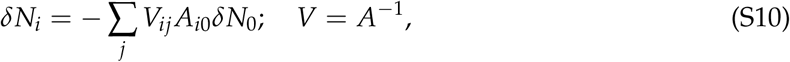

where *V*_*ij*_ is also the environmental sensitivity matrix ∂*N*_*i*_/∂*r*_*j*_ of the resident community. In other words, the invader acts as the following persistent perturbation of resident species growth parameters:

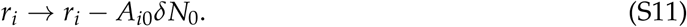

Taking the perspective of the invading species, its abundance *δN*_0_ should satisfy:

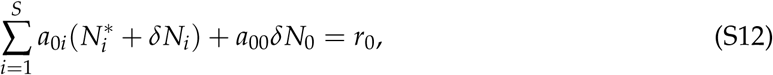

which, to first order in *δN*_0_ and *δN*_*i*_, becomes

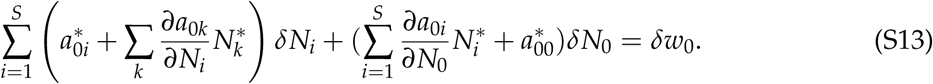

We define the interactions *A*_0*i*_ from resident species to invader so that *A*_0*i*_ encompasses the effect that species *i* can have on the other interactions that can affect the invader. The self-interaction term *A*_00_ encodes direct intra-specific interactions as well as self-induced changes of interactions with other species. With these notations we finally get

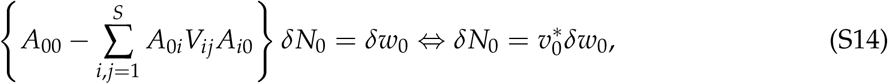

recovering the fitness-feedback factorization of the abundance of the invader. If the latter has finite (i.e., not infinitesimal) invasion fitness *w*_0_, if it coexists with resident species its abundance will read

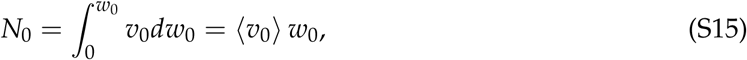

where ⟨*v*_0_⟩ is the average feedback over the fitness range [0, *w*_0_]. In general there will be no closed form for ⟨*v*_0_⟩. We can expect however, that it will be strongly correlated with its value 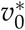 near the bifurcation *w*_0_ = 0. In particular, for Lotka–Volterra models 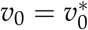.

#### S2.1 Feedback is caused by indirect effects

To illustrate the role that indirect effects play in the community feedback, we focus (for simplicity) on Lotka–Volterra models with self-regulation, i.e where all *a*_*ii*_ > 0. In this case we can define a general carrying capacity (i.e allowing negative values) *K*_*i*_ = *r*_*i*_/*a*_*ii*_ for all species. We then have a non dimensional feedback expression

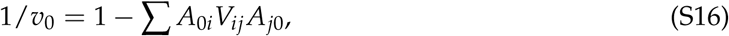

where *V*_*ij*_ = ∂*N*_*i*_/∂*K*_*j*_, and relative interactions *A*_0*i*_ = *a*_0*i*_/*a*_00_ and *A*_0*i*_ = *a*_*j*0_/*a*_*jj*_. Furthermore,

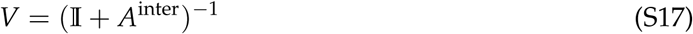

where 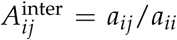 and 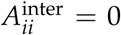 denotes relative *interspecific* interactions. 𝕀 is the identity matrix (*intraspecific* interactions). To avoid cumbersome notations, we will simply write *A* instead of *A*^inter^, remembering that its diagonal is now set to zero. We can then develop *V* in Neumann series

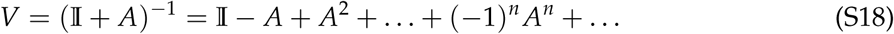

This means that

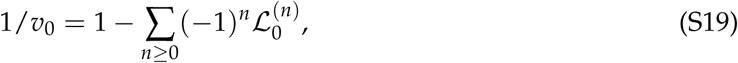

where 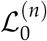 represents the sum of all invader-invader interaction loops of length *n* + 2 through the community:

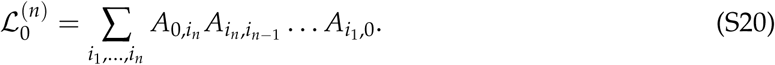

### S3 Analytical predictions for regime boundaries

#### S3.1 Two-species case

In the two species case treated in the main text,

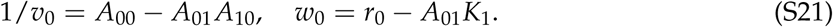

Our analysis in one of transcritical bifurcations. As parameters change, transition between invasion outcomes (Fig. 4a in main text) occur when equilibria meet and exchange their stability. For instance, the resident state (no invader) becomes unstable at (*w*_0_ = 0) which is where it crosses the coexistence state. On the other hand, the extinction of the resident occurs when the coexistence becomes unstable (and vice-versa) after crossing the invader state (no resident). Thus, when

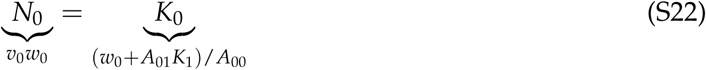

we get the condition

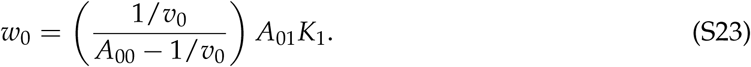

We see that for 1/*v*_0_ ≥ *a*_00_, there is no possible crossing, hence the invasion cannot cause the exclusion of the resident population. This boundary between domains which behaves as 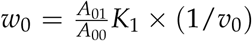 near 1/*v*_0_ = 0, and as −*A*_01_*K*_1_ for 1/*v*_0_ → −∞. The boundaries between regions in the fitness-feedback plane (shown in Fig. 4) are thus functions of carrying capacities and interactions, except for the 1/*v*_0_ = 0 and *w*_0_ = 0 boundaries which are universal.

To make the argument more general, it is useful to realize that the sensitivity of invasion fitness *w*_0_ to a change of abundance (*N*_1_) of the resident species is 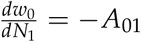.

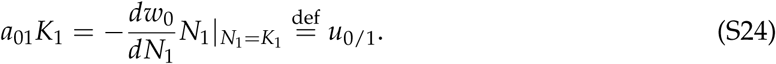

Furthermore 1/*a*_00_ is no other than the feedback *v*_0/1_ *in the absence of species 1*. With these notations the intersection between the coexistence state and the monospecific one becomes

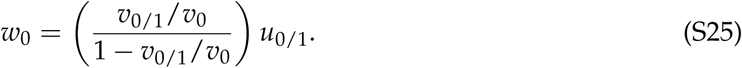

#### S3.2 General case

In the general multi-species case we can make a similar argument: the transition between invasion outcomes coincide with transcritical bifurcations which occurs when equilibria meet and exchange their stability (cf. Fig. S2). Assuming that there is at most one stable state per species composition, the coexistence state can become unstable (or vice versa) when it meets a state where one species, say *i*, is absent, i.e., when

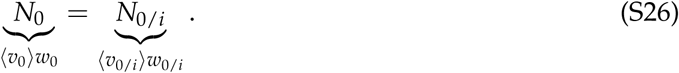

We focus on *w*_0/*i*_, the invasion fitness of species 0 in the absence of species *i*. We have

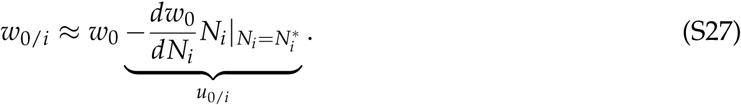

This expression is exact for the Lotka–Volterra case because fitness is a linear function of abundances. However, it will remain a good approximation in more general models when the first species to go extinct due to the invasion has a small abundance, thus justifying this linear approximation. The intersection condition thus reads

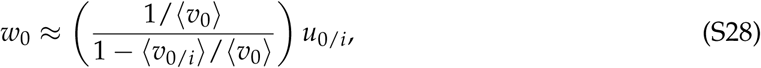

where we recognize a similar expression to the one derived in the two-species case. The vertical asymptote, that defines a bound on the turnover region in the fitness-feedback map is

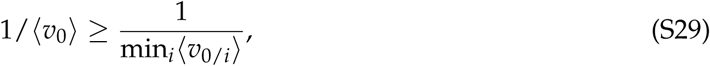

and the alternative state regime is limited to *v*_0_ < 0 and

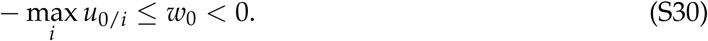

These bounds are not universally determined by ⟨*v*_0_⟩ and *w*_0_. The complex functions of dynamical parameters max_*i*_ *u*_0/*i*_ and min_*i*_⟨*v*_0/*i*_⟩ play an explicit role, and cannot be expected to be independent of ⟨*v*_0_⟩ and *w*_0_. This is why the transitions can be smooth in Fig. 4b of the main text.

**Figure S1:**
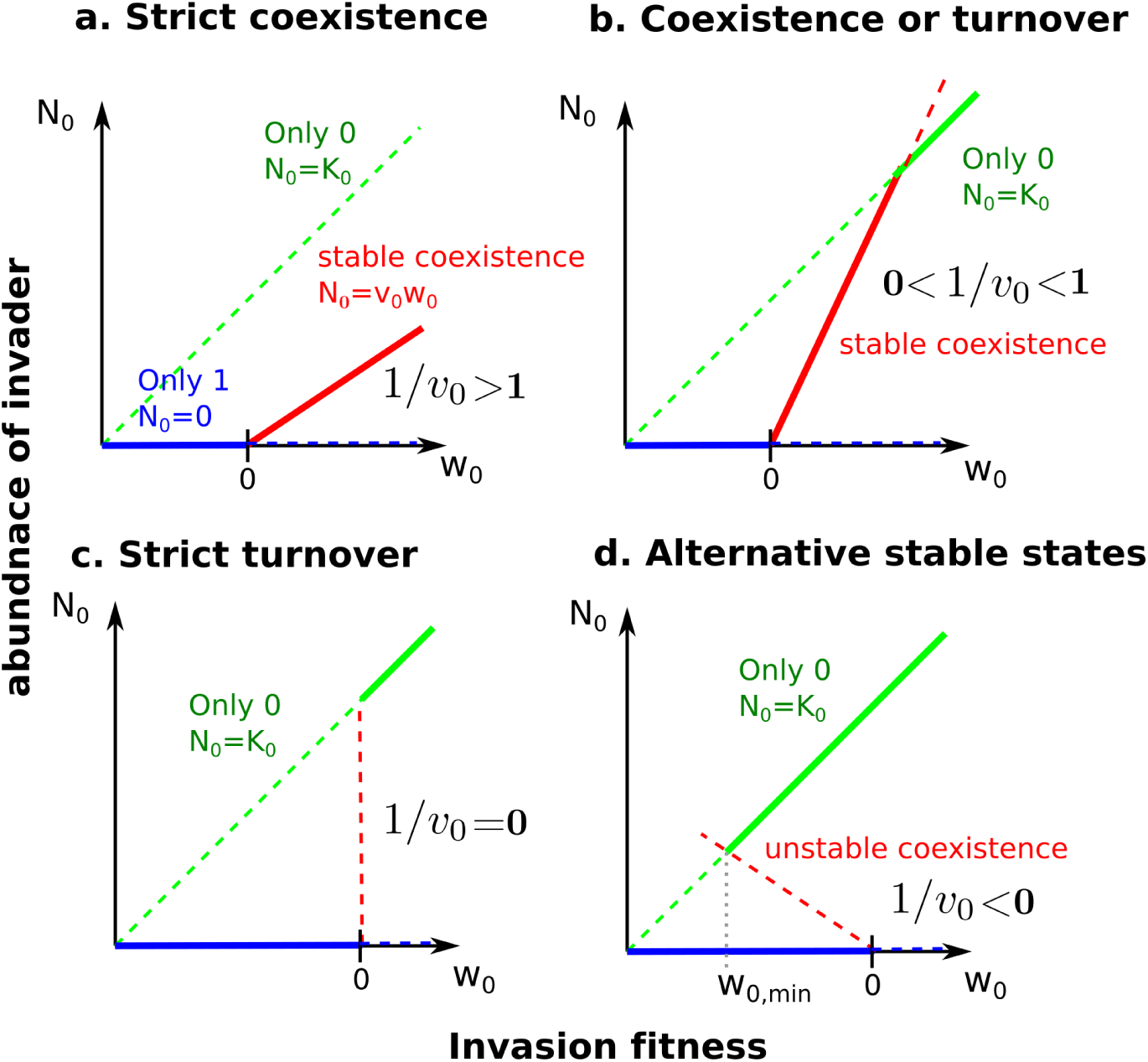
Bifurcation diagram of equilibrium abundance *N*_0_ of species 0 as a function of its invasion fitness *w*_0_ = *K*_0_ − *A*01 *K*1 into a resident population from species 1. Dashed lines represent unstable, and solid lines stable states. There are three possible states: (i) Species 1 persists alone, *N*_0_ = 0 (blue line). (ii) Species 0 replaces species 2, *N*_0_ = *K*_0_ (green line). (iii) Coexistence, *N*_0_ = *v*_0_ *w*_0_ (red line). If *w*_0_ < 0, species 1 cannot invade. The feedback strength *v*_0_ = ∂*N*_0_/∂*K*_0_ defined in (14) gives the slope of the graph of *N*_0_(*w*_0_) in the coexisting equilibrium. *v*_0_ allows to determine which of the three possible states will be reached following the invasion. (a,b) if 1/*v*_0_ > 0 species can coexist either (a) for all positive values of *w*_0_ or (b) only up to a threshold in *w*_0_ beyond which the resident species is displaced. (c,d) If 1/*v*_0_ ≤ 0, species exclude each other. (c) 1/*v*_0_ = 0 is the classical setting of adaptive dynamics, where a mutant phenotype fixes in the population as soon as its fitness is positive. (d) For 1/*v*_0_ < 0 the coexisting equilibrium is unstable and separates two alternate states (only species 0 survives, or only species 1). For *w*_0_ ∈ [*w*_0,min_, 0], these two states are both stable, meaning that species 0 will be able to invade only if its initial abundance is high enough (e.g., high propagule pressure is needed for invasion to take place).

**Figure S2:**
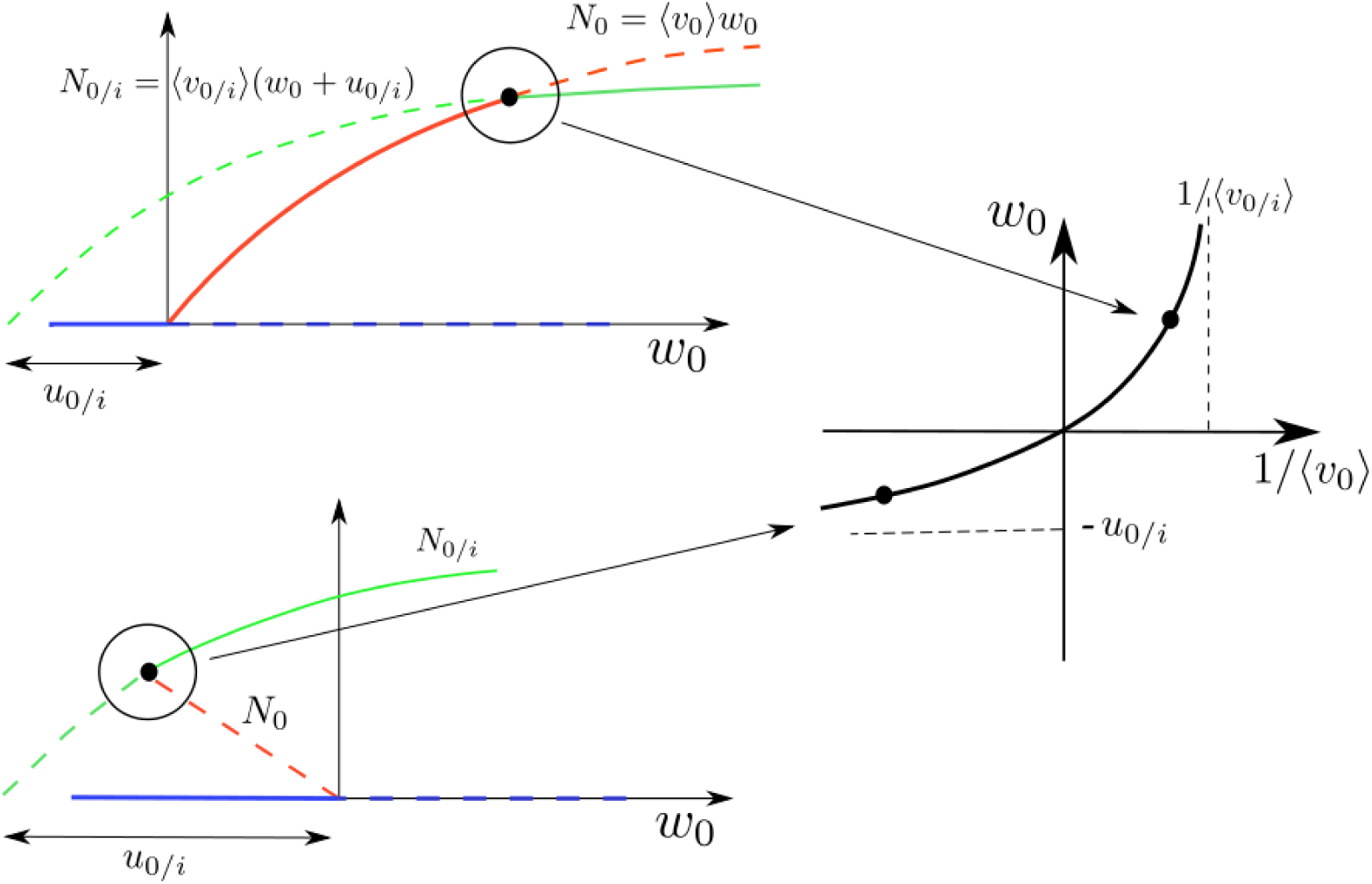
Illustration of the derivation of regime boundaries. Left panels: y-axis represents the possible equilibrium abundances of the invader as a function of its invasive fitness *w*_0_. Coloured dashed lines denote unstable states, continuous lines stable states. Top: the case where *v*_0_ > 0. As long as the invasion fitness is not too large, the coexistence state remains stable. As *w*_0_ increases, coexistence looses its stability when it crosses another state where the invader can persist (here a state where one species *i* has been led to extinction). If the feedback is small enough, however, the invader will not cross any other states, no matter how large its invasion fitness is (vertical asymptote on the right panel) Consequently, it will not cause extinctions. Bottom left, the case where *v*_0_ < 0. In this case the coexistence state is unstable, no matter how small the invasion fitness is, so that if the invasion fitness is positive, the community will shift to a different state. At negative values of invasion fitness, the unstable coexistence state delineates the basin of attraction of two alternative stable states: one where the invader is extinct (in blue) and one where it persists, at the expense of one (or more) species. At sufficiently negative values of *w*_0_ there is no stable state of the community in which the invader can persist (horizontal asymptote on the right panel).

### S4 Competitive exclusion in resource-consumer networks

Consider a bi-partite trophic network, where consumers (with indices *i, j*) feed on many resources (with indices *a, b*), following

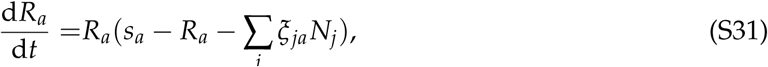

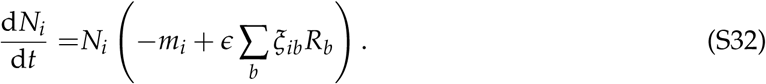

The equilibrium condition for resources is

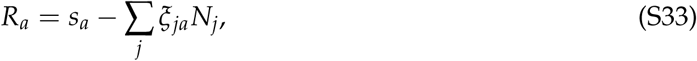

which can be inserted in the equilibrium condition for consumers:

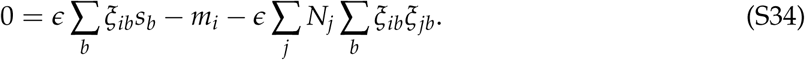

We thus find an effective Lotka–Volterra-like equilibrium for consumers:

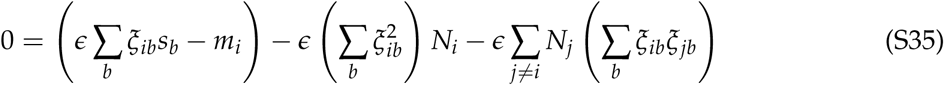

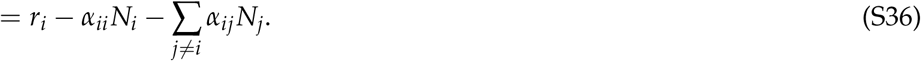

We can now perform our invasion analysis. We take the perspective of the consumers, whose abundances *N*_*i*_ satisfy

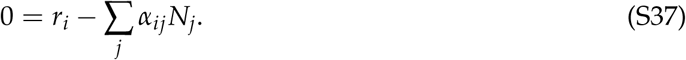

We can define the environmental sensitivity matrix *V* to encode the response of the community to a change of growth rate (rather than carrying capacity as in the main text and section S2). Thus,

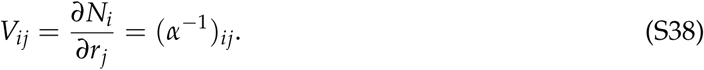

We add an invading species 0, whose interactions with the rest of the community are given by two vectors of coefficients, which we denote as follows:

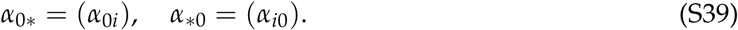

If the invader can coexist with all species present, its equilibrium will read

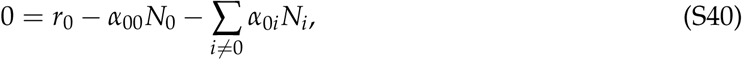

and as before, we assume a small effect of the invader on the existing community, so that

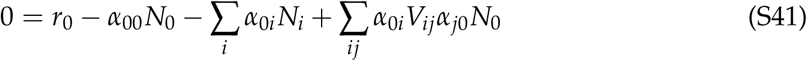

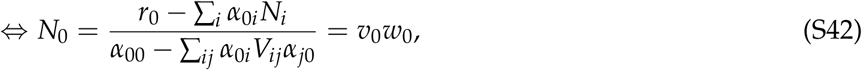

Where, in matrix notation,

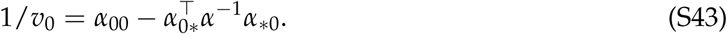

The interaction coefficients, however, have a very particular structure, inherited form the consumption coefficients *ξ*_*ia*_. Indeed, we have, if *ξ*_0_ = (*ξ*_0*a*_), that

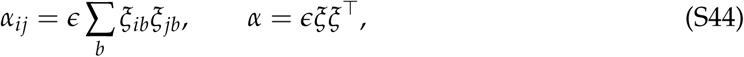

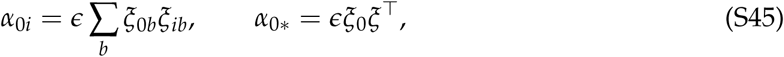

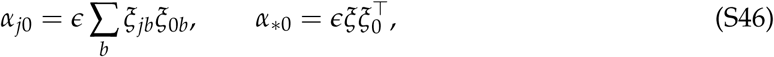

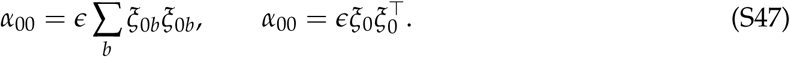

The consequence is that

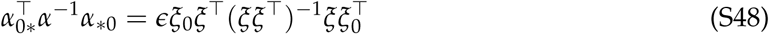

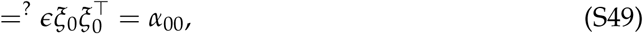

where the equality sign in (S49) is marked as a question, because going from the preceding line to this one is not so obvious: it requires all the involved matrices being invertible. This implies, in particular, that the number of surviving predators is equal to the number of preys, hence that the community is “saturated”. Assuming that (S49) holds, we find that

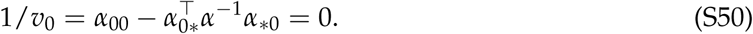

In other words, the structure of resource competition has a peculiar effect: it constrains the coexisting equilibrium to the limit case 1/*v*_0_ = 0, where a better competitor will always exclude a poorer one on the same resources, and mutual exclusion (in fact, any kind of multistability) is impossible. In this calculation, however, we used matrix notations in a fairly cavalier way. Imagine for instance that species 0 consumes only one resource that no other species is consuming. Then, *α*_0*i*_ = *α*_*i*0_ = 0 while *α*_00_ ≠ 0, so the previous calculation cannot hold (and *v*_0_ = 1/*α*_00_ is finite).

More generally, if there are more resources than species at the pre-invasion equilibrium *N*^∗^, then (*ξξ*^*⊤*^)^−1^ may have a nonzero kernel, meaning that line (S48) is in fact not equal to line (S49), but the corresponding sum is missing some terms. This corresponds to the case where invader 0 is somehow “innovating” by consuming an unused resource or mix of resources, in such a way that it potentially becomes an equal competitor to others in the community, rather than being excluded by another or excluding them.

### S5 v_0_ and the principle of limiting similarity

According to Eq. (8) in the main text, the invader’s equilibrium density 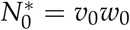 is the product of the invasion fitness *w*_0_ and the sensitivity *v*_0_, where

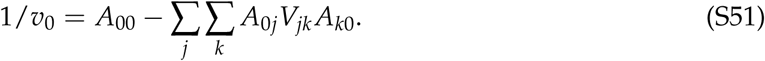

Here we show that if the invader has identical interaction coefficients to another species (let that be species 1 without loss of generality), then 1/*v*_0_ = 0 implying infinite sensitivity.

Indeed, using the fact that *V* = *A*^−1^ is the environmental sensitivity matrix (Eq. S10; see also Meszéna *et al.* 2006, Barabás *et al.* 2014), we can write 1/*v*_0_ as

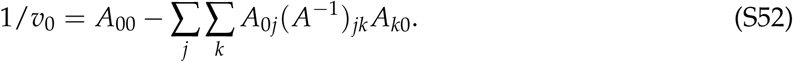

Since the invader (species 0) and species 1 have identical interaction coefficients by assumption, the zero subscripts can be replaced with ones:

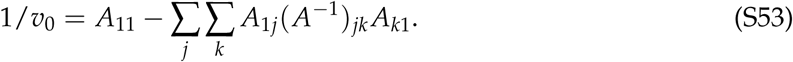

Focusing on just ∑_*k*_(*A*^−1^)_*jk*_ *A*_*k*1_, this is the product of the inverse of a matrix and the first column of the original matrix. The result, by definition, is a column vector with 1 as its first entry and 0s everywhere else. We therefore have

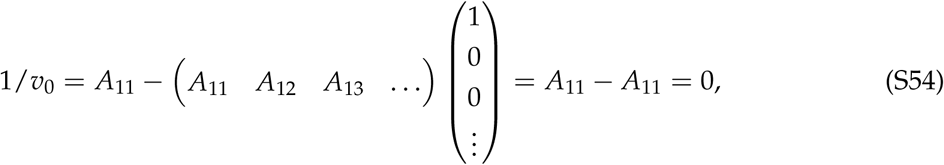

yielding infinite sensitivity.

Eq. (S54) was derived assuming the invader’s interactions *A*_0*i*_ and *A*_*i*0_ are identical to those of one of the resident species. Since 1/*v*_0_ is a continuous function of the matrix entries, for an invader with nonidentical but highly similar interaction structure to that of a resident, 1/*v*_0_ will be close to zero, giving a very large (though finite) sensitivity. Being sufficiently different from all residents is therefore a necessary condition for *v*_0_ not to be too large and thus for the approximation 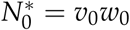 to work.

This result is in line with the theory of adaptive dynamics (Metz *et al.* 1996, Dieckmann & Law 1996, Geritz *et al.* 1998b, Meszéna 2005, Meszéna *et al.* 2005, Brännström *et al.* 2013), and our intuition of natural selection in general. If two species are exactly identical, then the slightest advantage to one of them (say, a mutation which reduces mortality by 0.1% on average) will cause it to establish and drive its competitor extinct. The fact that in the deterministic limit an arbitrarily small such parameter perturbation can change the identity of the winning species is reflected in the sensitivity *v*_0_ being infinitely large. In the terminology of adaptive dynamics, mutant phenotypes that are very similar to a resident one will either go extinct, or themselves become resident, replacing the original resident phenotype. The only exception is at an evolutionary branching point (Geritz *et al.* 1998b), where two competing phenotypes may start diverging from one another without competitive exclusion. However, the way such branching points permit the temporary stable coexistence of arbitrarily similar phenotypes is not by making *v*_0_ smaller, but having the fitness difference *w*_0_ between phenotypes be fine-tuned so that *v*_0_*w*_0_ will remain bounded thus coexistence, even when *v*_0_ is arbitrarily large.

In summary, an invading species that is very similar to a resident has very large sensitivity *v*_0_, rendering the approximation 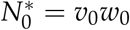 incompatible with the assumption of small 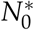. This is because such a “mutant” phenotype either goes extinct or replaces the original resident. Neither scenario conforms to the assumption of the invader establishing at a low density and affecting the resident community only slightly. Whether the mutant invader will replace the resident can be determined via the methods of adaptive dynamics; e.g., by comparing their invasion growth rates. The only exception to this rule of competitive exclusion is when the dynamics are exactly at an evolutionary branching point, where the special parameter settings required to offset the effect of a large *v*_0_ are automatically established.

### S6 Interaction between community stability and invader properties

#### S6.1 Worst-case invader

A resident community in which the environmental sensitivity matrix *V* has large elements is more likely to allow some invading species to experience positive indirect feedback. In the main text we formalized this intuition by defining the worst-case bound

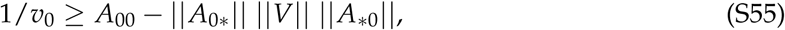

where ‖ · ‖ is the Euclidean norm for interaction vectors *A*_0∗_ = (*A*_0*j*_), *A*_∗0_ = (*A*_*i*0_), and the spectral norm for the environmental sensitivity matrix *V* (the max of ‖*Vu*‖ over all normalized vectors *u*). The larger the norms ‖*A*_0∗_‖ and ‖*A*_∗0_‖ the larger the feedbacks but the latter also involves the environmental sensitivity matrix *V*. Here we give a method to compute the worst-case invader. We start from the expression of the spectral norm of *V*:

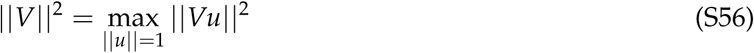

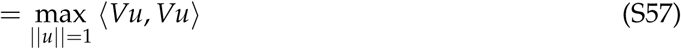

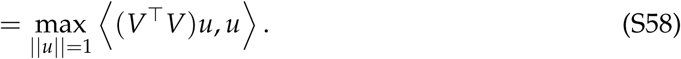

From this expression we deduce that the max is obtained by choosing *u* = *u*_dom_ where *u*_dom_ is the (normalized) eigenvector associated with the dominant (i.e largest) eigenvalue 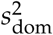 of the positive definite matrix *V*^*⊤*^*V*. We also deduce that ‖*V*‖ = *s*_dom_ (which is why this norm is called “spectral”). To elicit the largest long-term impact, the invader-community vector *A*_∗0_ must align with the eigenvector *u*_dom_. Thus,

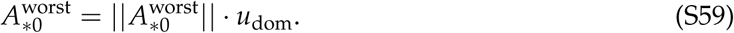

Then, the community-invader vector *A*_∗0_ must align with the direction of the associated community response *Vu*_dom_:

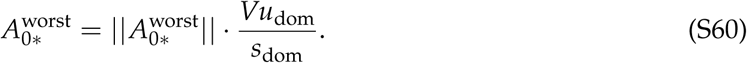

#### S6.2 Mean-case invader

We now seek the expected value of the indirect feedback on a species randomly chosen from a pool of potential invaders. This requires prior knowledge on the first moments *M*_*ij*_ = 𝔼(*A*_*i*0_ *A*_0*j*_) and *d* = 𝔼(*A*_00_) of a statistical ensemble of invader-community interactions. Indeed, the long-term feedbacks on an invader reads

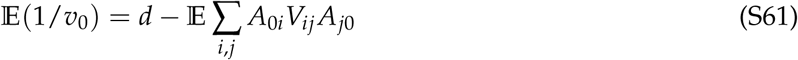

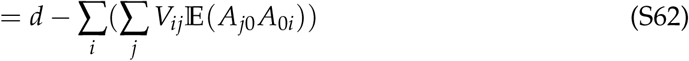

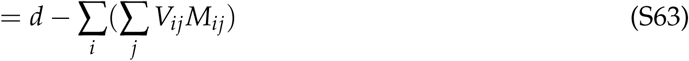

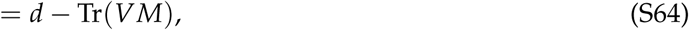

where Tr(*B*) stands for the trace of a given matrix *B*. In the main text we chose a particularly simple pool of invaders, corresponding to

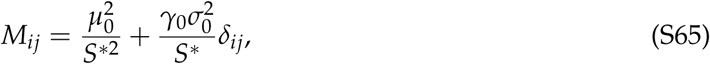

and *d* = 1. We also considered the hight diversity limit *S*^∗^ → ∞. From the above calculations we get

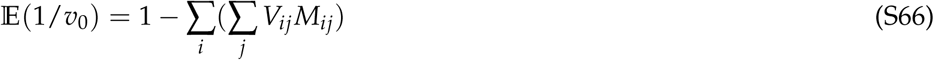

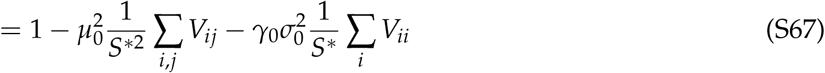

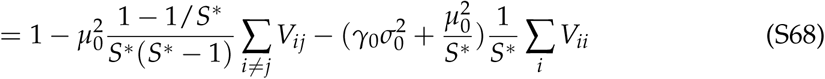

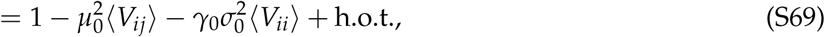

where “h.o.t.” stands for “higher-order terms” in 1/*S*^∗^, that vanish at high diversity.

### S7 Many-species simulations

In the main text, numerical results shown in Fig. 4 and 5 were obtained from the Lotka–Volterra model (or a nonlinear extension, see below), whose dynamics are given by

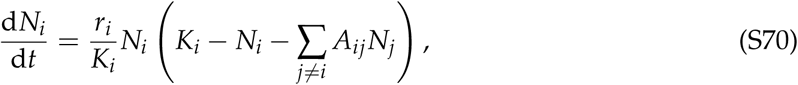

with *r*_*i*_ the growth rate of species (or strategy) *i*. For simplicity, we assume *r*_*i*_ = *K*_*i*_. This model was chosen as a simple test for our analysis, and we discuss its specificities in the following section. We now describe the procedure for generating the numerical results shown in the text.

#### S7.1 Disordered LV communities: Generating Figures 4b and 5 (upper panel)

We generate a random community of *S* = 10 species with normally distributed, symmetric interactions *a*_*ij*_ = *a*_*ji*_ with statistics 𝔼(*a*_*ij*_) = 0.5, std(*a*_*ij*_) = 0.25, and all carrying capacities *K*_*i*_ = 1. We simulate the dynamics defined by Eq. (S70) and measure the number of survivors *S*^∗^ (species with abundances above *N*_*c*_ = 10^−10^ at time 10^5^) and their equilibrium abundances 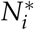.

We then draw 500 random invaders with *K*_0_ = 1 and symmetric interactions *a*_0*i*_ = *a*_*i*0_, drawn from a normal distribution. The statistics of this distribution are chosen to ensure that we get both positive and negative values of *w*_0_ and 1/*v*_0_:

- 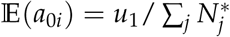 with *u*_1_ a uniform random number drawn in the interval [0.7, 1.3]
- 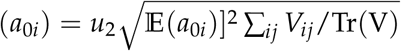 with *u*_2_ drawn uniformly in the interval [0, 2].

For each invader independently, we add it to the original community equilibrium and simulate its dynamics from two initial conditions: 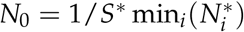 and 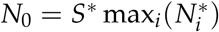, to test for multistability.

**Figure S3:**
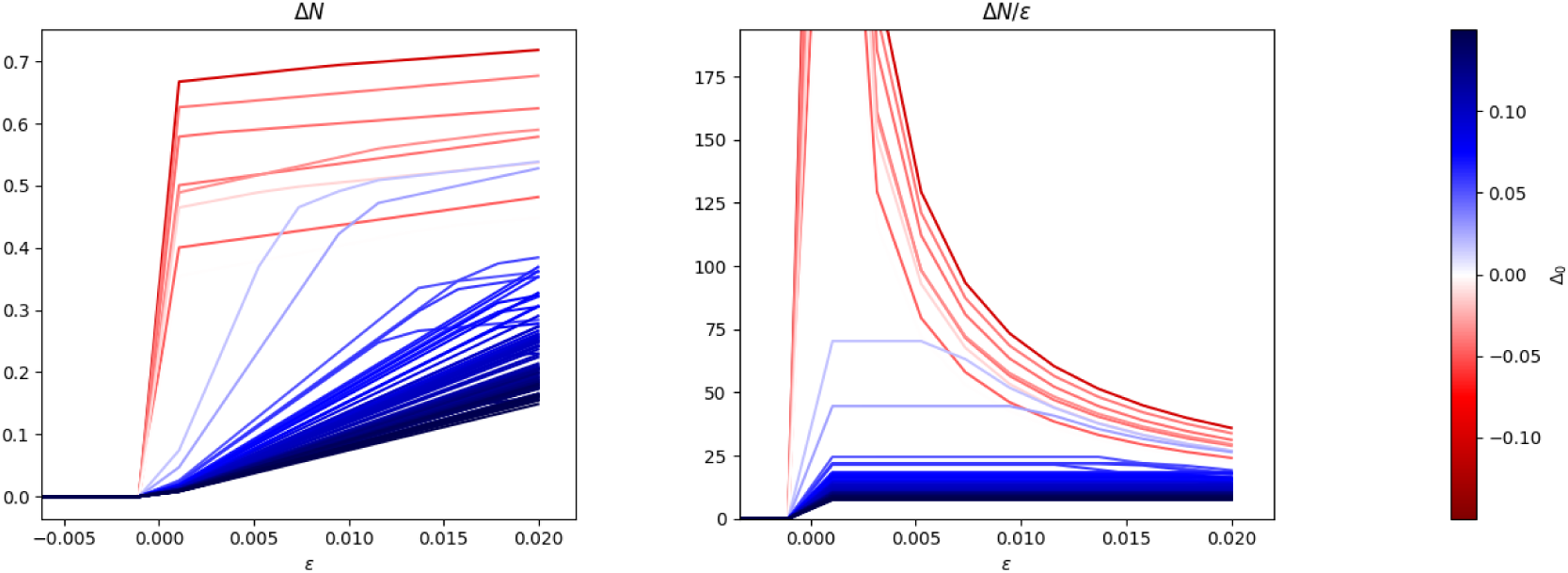
Change in equilibrium composition 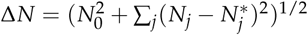 after the invasion of species 0, as a function of *w*_0_ = *ϵ* the invasion fitness. Each line corresponds to a randomly drawn invader with Δ_0_ = 1/*v*_0_ shown by colors. **Left:** We see that for reversible invaders (blue, *v*_0_ > 0) the equilibrium starts to change smoothly at the invasion threshold *w*_0_ = 0, whereas for irreversible invaders (red, *v*_0_ < 0) there is a sudden jump. **Right:** For reversible invaders Δ*N*/*w*_0_ tends to a constant when *w*_0_ → 0, reflecting the fact that the change in community is smooth and proportional to *w*_0_, while it diverges for irreversible invaders.

We then measure the number of extinctions as a result of invasion, the existence of alternate states, and the values of *v*_0_ and *w*_0_ for each invader, and use these properties to locate each invader on our map (Fig. 4) and label them with the different categories: no invasion, coexistence, turnover, irreversible turnover, and alternative stable states. The color background of Fig. 4 is obtained by training a Support Vector Classifier on these labelled data, predicting the label for each point in a grid covering the (1/*v*_0_, *w*_0_) plane, and assembling these predictions in a contour plot, using Python and the scikit-learn library.

#### S7.2 Two level Food-Webs: Generating Figures 4d and 5 (lower panel)

For the food webs considered in Fig. 4c-d, we first draw niche positions *n*_*i*_ ∈ [1, 2, 3] at random for *S* = 20 species. We then generate the matrix of predator-prey fluxes with randomly distributed coefficients (following the uniform distribution 𝒰)

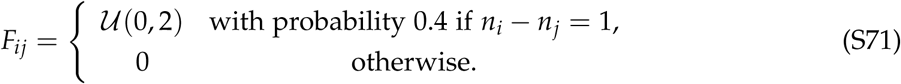

We then simulate the dynamics

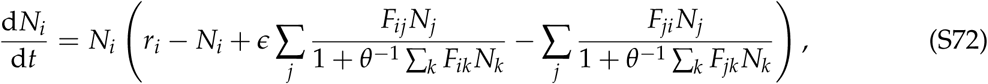

with *ϵ* = 0.4, *θ* = 0.1, and intrinsic growth rate

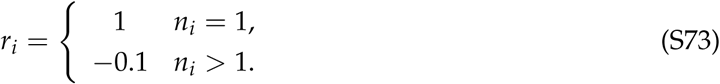

Invaders were not drawn according to the same structure, but with direct non-trophic interactions as in the previous section, so as to be able to visit the plane (1/*v*_0_, *w*_0_) in a less constrained way. Since interactions were indiscriminate of trophic level, they were often relatively strong against low-abundance predators, and all successful invasions caused species extinctions in the community, so we set the threshold for blue dots (“coexistence”) to less than 10% extinctions. Despite this quantitative difference, the same qualitative pattern held as in random Lotka–Volterra communities.

## Footnotes

a If *w*_0_ lies under the curve (16) or its many-species extension in Appendix S3.

1 In Appendix S1 we show that a similar expression can be extended to any value of invasion fitness.

2 If *v*_0_ < 0, the abundance 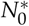 given by (6) is negative, meaning that the weak-impact approximation is incorrect, however small the invasion fitness *w*_0_ may be. The coexistence state is unstable, and the invasion threshold *w*_0_ = 0 is a tipping point.

3 Notice the log scale in the y-axis of Fig. 5.

4 The maximum of ‖*Vu*‖ over all normalized vectors *u*.

5 This worst-case a special case. Yet we show in Appendix S6.2 that an expression analogous to the worst-case (17) can be derived for a mean-case scenario over a pool of potential invaders.

## Literature Cited

Anic, V., Henríquez, C., Abades, S.R. & Bustamante, R. (2015). Number of conspecifics and reproduction in the invasive plant eschscholzia californica (papaveraceae): is there a pollinator-mediated allee effect? Plant Biology, 17, 720–727.

Arnoldi, J.F., Loreau, M. & Haegeman, B. (2016). Resilience, reactivity and variability: A mathematical comparison of ecological stability measures. J. Theor. Biol., 389, 47–59.

Aufderheide, H., Rudolf, L., Gross, T. & Lafferty, K.D. (2013). How to predict community responses to perturbations in the face of imperfect knowledge and network complexity. Proceedings of the Royal Society of London Series B, 280, 20132355.

Barabás, G., D’Andrea, R. & Stump, S.M. (2018). Chesson’s coexistence theory. Ecological Monographs, 88, 277–303.

Barabás, G., Pásztor, L., Meszéna, G. & Ostling, A. (2014). Sensitivity analysis of coexistence in ecological communities: theory and application. Ecology Letters, 17, 1479–1494.

Barbier, M., Arnoldi, J.F., Bunin, G. & Loreau, M. (2018). Generic assembly patterns in complex ecological communities. Proceedings of the National Academy of Sciences, 115, 2156–2161.

Blackburn, T.M., Pysek, P., Bacher, S., Carlton, J.T., Duncan, R.P., Jarosik, V., Wilson, J.R.U. & Richardson, D.M. (2011). A proposed unified framework for biological invasions. Trends in Ecology & Evolution, 26, 333 – 339.

Brännström, Å., Johansson, J. & Von Festenberg, N. (2013). The hitchhiker’s guide to adaptive dynamics. Games, 4, 304–328.

Bunin, G. (2018). Directionality and community-level selection. bioRxiv, p. 484576.

Catford, J.A., Smith, A.L., Wragg, P.D., Clark, A.T., Kosmala, M., Cavender-Bares, J., Reich, P.B. & Tilman, D. (2019). Traits linked with species invasiveness and community invasibility vary with time, stage and indicator of invasion in a long-term grassland experiment. Ecology Letters, 22, 593–604.

Champagnat, N., Ferriere, R. & Ben Arous, G. (2002). The canonical equation of adaptive dynamics: a mathematical view. Selection, 2, 73–83.

Chesson, P. (2000). Mechanisms of maintenance of species diversity. Annual review of Ecology and Systematics, 31, 343–366.

Courchamp, F., Berec, L. & Gascoigne, J. (2008). Allee effects in ecology and conservation. Oxford University Press.

Courchamp, F., Clutton-Brock, T. & Grenfell, B. (1999). Inverse density dependence and the allee effect. Trends in ecology & evolution, 14, 405–410.

Cuddington, K., Wilson, W., Hastings, A., Roelke, A.E.D.L. & DeAngelis, E.D.L. (2009). Ecosystem engineers: Feedback and population dynamics. The American Naturalist, 173, 488–498.

Dakos, V. (2018). Identifying best-indicator species for abrupt transitions in multispecies communities. Ecological indicators, 94, 494–502.

Dieckmann, U. & Law, R. (1996). The dynamical theory of coevolution: a derivation from stochastic ecological processes. Journal of Mathematical Biology, 34, 579–612.

Doebeli, M. & Dieckmann, U. (2000). Evolutionary branching and sympatric speciation caused by different types of ecological interactions. The american naturalist, 156, S77–S101.

Donohue, I., Hillebrand, H., Montoya, J.M., Petchey, O.L., Pimm, S.L., Fowler, M.S. et al. (2016). Navigating the complexity of ecological stability. Ecol. Lett., 19, 1172–1185.

Eisenhauer, N., Schulz, W., Scheu, S. & Jousset, A. (2013). Niche dimensionality links biodiversity and invasibility of microbial communities. Functional Ecology, 27, 282–288.

Elton, C.S. (1958). The ecology of invasions by animals and plants. The ecology of invasions by animals and plants.

Frost, C.M., Allen, W.J., Courchamp, F., Jeschke, J.M., Saul, W.C. & Wardle, D.A. (2019). Using network theory to understand and predict biological invasions. Trends in ecology & evolution.

Fukami, T. & Nakajima, M. (2011). Community assembly: alternative stable states or alternative transient states? Ecology letters, 14, 973–984.

Gaertner, M., Biggs, R., Te Beest, M., Hui, C., Molofsky, J. & Richardson, D.M. (2014). Invasive plants as drivers of regime shifts: identifying high-priority invaders that alter feedback relationships. Diversity and Distributions, 20, 733–744.

Galiana, N., Lurgi, M., Montoya, J.M. & López, B.C. (2014). Invasions cause biodiversity loss and community simplification in vertebrate food webs. Oikos, 123, 721–728.

Geritz, S.a.H., Kisdi, E., Meszéna, G. & Metz, J.a.J. (1998a). Evolutionarily singular strategies and the adaptive growth and branching of the evolutionary tree. Evolutionary Ecology, 12, 35–57.

Geritz, S.A.H., Kisdi, É., Meszéna, G. & Metz, J.A.J. (1998b). Evolutionary singular strategies and the adaptive growth and branching of evolutionary trees. Evolutionary Ecology, 12, 35–57.

Gilpin, M.E. & Case, T.J. (1976). Multiple domains of attraction in competition communities. Nature, 261, 40.

Grainger, T.N., Levine, J.M. & Gilbert, B. (2019). The invasion criterion: A common currency for ecological research. Trends in ecology & evolution.

Guo, Q., Fei, S., Dukes, J.S., Oswalt, C.M., III, B.V.I. & Potter, K.M. (2015). A unified approach for quantifying invasibility and degree of invasion. Ecology, 96, 2613–2621.

Hui, C. & Richardson, D.M. (2018). How to invade an ecological network. Trends in ecology & evolution.

Ives, A.R. & Carpenter, S.R. (2007). Stability and diversity of ecosystems. Science, 317, 58–62.

Jiménez-Valverde, A., Peterson, A.T., Soberón, J., Overton, J.M., Aragón, P. & Lobo, J.M. (2011). Use of niche models in invasive species risk assessments. Biological Invasions, 13, 2785–2797.

Kotil, S.E. & Vetsigian, K. (2018). Emergence of evolutionarily stable communities through eco-evolutionary tunnelling. Nature ecology & evolution, 2, 1644.

Kotta, J., Wernberg, T., Jänes, H., Kotta, I., Nurkse, K., Pärnoja, M. & Orav-Kotta, H. (2018). Novel crab predator causes marine ecosystem regime shift. Scientific reports, 8, 4956.

Kramer, A.M. & Drake, J.M. (2010). Experimental demonstration of population extinction due to a predator-driven allee effect. Journal of Animal Ecology, 79, 633–639.

Law, R. & Morton, R.D. (1996). Permanence and the assembly of ecological communities. Ecology, 77, 762–775.

Levine, J.M., Vila, M., Antonio, C.M.D., Dukes, J.S., Grigulis, K. & Lavorel, S. (2003). Mechanisms underlying the impacts of exotic plant invasions. Proceedings of the Royal Society of London. Series B: Biological Sciences, 270, 775–781.

Levins, R. (1974). Qualitative analysis of partially specified systems. Ann. NY Acad. Sci., 231, 123–138.

Lewis, M.A., Petrovskii, S.V. & Potts, J.R. (2016). The mathematics behind biological invasions. vol. 44. Springer.

Liautaud, K., van Nes, E.H., Barbier, M., Scheffer, M. & Loreau, M. (2019). Superorganisms or loose collections of species? a unifying theory of community patterns along environmental gradients. Ecology letters.

Lurgi, M., Galiana, N., López, B.C., Joppa, L.N. & Montoya, J.M. (2014). Network complexity and species traits mediate the effects of biological invasions on dynamic food webs. Frontiers in Ecology and Evolution, 2, 36.

MacDougall, A.S., Gilbert, B. & Levine, J.M. (2009). Plant invasions and the niche. Journal of Ecology, 97, 609–615.

McLellan, B.N., Serrouya, R., Wittmer, H.U. & Boutin, S. (2010). Predator-mediated allee effects in multi-prey systems. Ecology, 91, 286–292.

McNally, L. & Jackson, A.L. (2013). Cooperation creates selection for tactical deception. Proceedings of the Royal Society B: Biological Sciences, 280, 20130699.

Meszéna, G. (2005). Adaptive dynamics: the continuity argument. Journal of Evolutionary Biology, 18, 1182–1185.

Meszéna, G., Gyllenberg, M., Jacobs, F.J. & Metz, J.A.J. (2005). Link between population dynamics and dynamics of darwinian evolution. Physical Review Letters, 95, 078105.

Meszéna, G., Gyllenberg, M., Pásztor, L. & Metz, J.A.J. (2006). Competitive exclusion and limiting similarity: a unified theory. Theoretical Population Biology, 69, 68–87.

Metz, J., Geritz, S., Meszena, G., Jacobs, F. & van Heerwaarden, J. (1995). Adaptive dynamics: A geometrical study of the consequences of nearly faithful reproduction. Tech. rep., International Institute for Applied Systems Analysis.

Metz, J.A.J., Geritz, S.A.H., Meszéna, G., Jacobs, F.J.A. & van Heerwaarden, J.S. (1996). Adaptive dynamics: A geometrical study of the consequences of nearly faithful reproduction. In: Stochastic and spatial structures of dynamical systems (eds. van Strien, S.J. & Verduyn Lunel, S.M.). Proceedings of the Royal Dutch Academy of Science, Amsterdam, The Netherlands, pp. 183–231.

Metz, J.A.J., Nisbet, R.M. & Geritz, S.A.H. (1992). How should we define ‘fitness’ for general ecological scenarios? Trends in Ecology & Evolution, 7, 198–202.

O’Sullivan, J.D., Knell, R.J. & Rossberg, A.G. (2018). Metacommunity-scale biodiversity regulation and the self-organised emergence of macroecological patterns. Ecology Letters.

Pimm, S.L. (1991). The Balance of Nature? Ecological Issues in the Conservation of Species and Communities. University of Chicago Press.

Rossberg, A.G. & Barabás, G. (2019). How carefully executed network theory informs invasion ecology. Trends in ecology & evolution, 34, 385–386.

Roughgarden, J. (1983). Competition and theory in community ecology. The American Naturalist, 122, 583–601.

Scheffer, M., Carpenter, S., Foley, J.A., Folke, C. & Walker, B. (2001). Catastrophic shifts in ecosystems. Nature, 413, 591.

Schreiber, S.J. (2000). Criteria for Cr robust permanence. Journal of Differential Equations, 162, 400–426.

Taylor, C.M. & Hastings, A. (2005). Allee effects in biological invasions. Ecology Letters, 8, 895–908.

Tilman, D. (1982). Resource competition and community structure. Princeton university press.

Turelli, M. (1978). A reexamination of stability in randomly varying versus deterministic environments with comments on the stochastic theory of limiting similarity. Theoretical Population Biology, 13, 244–267.

Venkateswaran, V.R. & Gokhale, C.S. (2019). Evolutionary dynamics of complex multiple games. Proceedings of the Royal Society B, 286, 20190900.

White, E.M., Wilson, J.C. & Clarke, A.R. (2006). Biotic indirect effects: a neglected concept in invasion biology. Diversity and distributions, 12, 443–455.

Williamson, M. (1999). Invasions. Ecography, 22, 5–12.

Wright, J.P. & Jones, C.G. (2006). The Concept of Organisms as Ecosystem Engineers Ten Years On: Progress, Limitations, and Challenges. BioScience, 56, 203–209.

Yodzis, P. (1988). The indeterminacy of ecological interactions as perceived through perturbation experiments. Ecology, 69, 508–515.

